# The neural dynamics of familiarity representation during cued recall

**DOI:** 10.64898/2026.09.21.753245

**Authors:** Yeliz Dinç, Yang Shi, Gyula Kovács

**Affiliations:** Department of Biological Psychology and Cognitive Neurosciences, Institute of Psychology, Friedrich Schiller University Jena, D-07743 Jena, Germany

**Keywords:** face familiarity, imagery, electroencephalography, multivariate decoding, temporal generalization, cued recall

## Abstract

Face familiarity elicits reliable neural signatures during perception, yet how it is represented when faces are recalled from memory by using visual imagery and how these signals relate to perceptual representations, remains unclear. We combined perceptual and imagery conditions with electroencephalography and multivariate pattern analysis. In a perceptual session we presented of four famous individuals, while in the imagery session we used a cued recall paradigm and presented abstract symbols, associated to them. By using time-resolved multivariate decoding we tested if familiarity information is shared across stimulus categories, and if perceptual face familiarity representations overlap with processing during cued recall, related to imagery. The face perception condition showed typical and robust familiarity information over occipito-temporal electrodes. The symbolic cues also carried similar familiarity information, demonstrating cross-category familiarity processing between symbols and faces. During cued recall, familiarity information emerged over posterior regions and evolved into a prolonged and late information component over right anterior electrodes. Importantly, cross-domain temporal generalization analysis revealed an overlap between perceptual face familiarity patterns and recall/imagery related processing: perceptual face familiarity information mapped onto later processing during cued recall, with a pronounced and prolonged right anterior dominance and a shorter overlap over left posterior channels. Together, these results suggest that familiarity processing, similarly to category representations, is supported by overlapping neural patterns across perception and recall/imagery, transitioning from dynamic posterior representations, consistent with perceptual analysis, to a stable and prolonged representation over right anterior sites, that is compatible with a sustained processing, related to recall and/or imagery.

## Introduction

In everyday life we are constantly exposed to different faces, which share numerous common features. Yet, we can effortlessly recognize familiar faces among unfamiliar individuals, even though people’s faces are constantly undergoing substantial changes, for example due to aging or by merely getting a haircut. This ability serves important social functions, such as being able to build lasting relationships or recognizing people. In addition to our ability of recognizing familiar people, we can also recall the faces of people we know, using our “mind’s eye”. The recall of someone’s face from memory is often elicited by cues that are either direct, such as a name or voice, or abstract and symbolic such as the logo of the company where someone is employed. While there is a large literature covering the neural processing of perceptual face familiarity using both uni and multivariate neuroimaging and electrophysiological methods (Ambrus et al., 2021; Dalski et al., 2022; Dubois et al., 1999; Gobbini & Haxby, 2006; Kovács, 2020; Li et al., 2022; Schweinberger et al., 2002; Shah et al., 2001; Wiese et al., 2022; for review, see Wiese et al., 2024) the neural mechanisms of imagery recall of familiar faces, especially of those cued by abstract stimuli or concepts, remains relatively understudied and unclear.

Cued recall includes the vivid mental imagery of the person as well as the actual retrieval of the person’s face from memory. Previous neuroimaging studies have demonstrated that visual perception and imagery share their underlying neural mechanisms (Dijkstra et al., 2018; Ragni et al., 2020, 2021; Xie et al., 2020). For example, similar to perceived stimuli, category and familiarity information can also be decoded for visually imagined stimuli from the fMRI activity pattern in a wide range of regions, including early visual, inferotemporal, parietal, and prefrontal regions (Ragni et al., 2021). Earlier univariate fMRI work showed that imagery of faces and places recruits corresponding posterior category-selective visual regions that are also involved in perception (O’Craven & Kanwisher, 2000). Moreover, distinct subdivisions of the medial parietal cortex were shown to be recruited during recall of familiar people and places while these regions respond negatively, but still in a category-selective manner, to visual face and scene presentations (Silson et al., 2019a). Furthermore, Xie et al. (2020) have shown, using EEG and multivariate pattern analysis (MVPA), that perceived and imagined stimuli share neural representations over the parieto-occipital cortex in the alpha EEG band (for a different conclusion see Shatek et al., 2019). However, it seems that the temporal dynamics of perceived and imagined stimuli are not identical. In visual perception, there is a well-known hierarchical processing stream, starting with early visual areas and spreading towards higher-level, more anterior brain regions. On the other hand, information during visual imagery becomes apparent only later, presumably due to slower developing, top-down processes. Importantly, the representation of imagined stimuli is similar to the higher-level processing stages of perceived stimuli, suggesting that imagery directly activates high-level visual areas (Dijkstra et al., 2017, 2018, 2019).

Regarding cued recall, intracranial recordings suggest that the hippocampus acts as a switchboard between perception and memory: hippocampal higher gamma band power increases after successful recall in about 500 ms after the cues. This, in turn, is followed by a change in the representation from cue to target-related information over extrahippocampal electrodes (Treder et al., 2021). Recently, two studies investigated the cued recall of familiar places and people. Corriveau et al. (2023) used MEG and a retro cue paradigm in which two written names were presented before a numerical cue indicated which personally familiar person or place should be recalled during a 4 second recall period. Category level representations emerged approximately 1000 ms after the retro cue and remained stable and temporally generalizable throughout the recall period. In contrast, Scrivener et al. (2025) used a single 500 ms verbal cue followed directly by a 2500 ms imagery period and combined EEG with fMRI-based representational fusion. People vs places category information was the strongest during the first 700 ms of imagery, followed by weaker sustained category information, while information distinguishing individual imagined stimuli re-emerged later, at approximately 700 ms for places and 1100 ms for people. Differences in cueing structure, task timing, and methodological approach may have contributed to the different temporal profiles observed across these two studies.

In the current study, we tested the recall of familiar persons, cued by associated symbols. Our primary objectives were the following. First, we aimed to examine the temporal dynamics of familiarity processing in the brain, elicited by the cued recall of famous faces. Second, we tested if familiarity information can also be observed for graphical symbols of abstract concepts, similarly to faces (Dalski et al., 2022; Gobbini & Haxby, 2006; Li et al., 2022), scenes (Klink et al., 2023) and other perceptual stimulus categories (Ely et al., 2026). Third, we tried to get a better understanding of how stable such familiarity representations are over time. Finally, using cross-domain decoding methods, we tested if the neural representations during familiar person perception overlap with those of the recall phases.

We applied a cued recall paradigm in which participants were presented with symbols representing famous films (Pirates of the Caribbean, Iron Man, Harry Potter) or with the national flag of the USA (and related symbols) as familiar and four, visually similar but unfamiliar symbols as control stimuli. By seeing these symbols, participants were instructed to recall and imagine the faces of the best corresponding associated person’s (Johnny Depp, Robert Downey Jr., Daniel Radcliffe and Donald Trump) while they underwent EEG recording procedures. In a separate, “localizer” paradigm, at the end of the experimental session, they were also presented with naturally variable images of the associated persons’ faces, as well as with faces of four unfamiliar male persons. Using time-resolved decoding, we tested the temporal dynamics of familiarity representations during the cued recall phase, elicited by symbolic abstract stimuli.

## Materials and Methods

### Participants

We recruited 37 participants. The data of 34 participants were analyzed (5 males, mean age = 22.52 years, SD = 3.62), as three of them had incomplete EEG recordings. Participants received partial course credits upon participation. This sample size is comparable to that of previous studies investigating the neural correlates of personal face and scene familiarity (Wiese et al., 2019; Ambrus et al., 2021; Li et al., 2022; Klink et al., 2023). A power analysis (G*Power; Faul et al., 2007), based on the lowest effect size of the cross-category decoding results of Klink et al., (2023) confirmed that this sample size is larger than the one necessary to obtain a power of 0.99 (paired-sample t-test; D = 1.2; n = 15). Participants were right-handed except three of them and had normal or corrected-to-normal vision without any history of neurological disorders. The study was approved by the ethics committee of the Friedrich Schiller University of Jena, and informed consents were obtained upon participation.

### Stimuli

Stimuli were acquired from the publicly available domain of the worldwide web and consisted of the faces of eight persons, as well as eight graphical symbols, each with 5 different images. Four of the identities were very well-known male celebrities (Johnny Depp, Robert Downey Jr., Daniel Radcliffe, and Donald Trump), related to the chosen symbols, validated in a separate behavioral pilot study (N = 16; see Supplementary Fig. 2). The other four identities were celebrities outside Germany (Goran Bogdan, Jorge Drexler, Alban Skënderaj, Uğurkan Erez), who were unfamiliar to our German participants (see Supplementary Materials). The graphical symbols, related to the familiar identities were selected from the most famous feature films of the actors (Pirates of the Caribbean, Iron Man, Harry Potter) as well as the national flag of the USA (and related symbols). As unfamiliar graphical symbols, we selected visually similar, complex company logos that were unrelated to any of the identities. None of these unfamiliar symbols were recognized by the participants in the pilot study (see Supplementary Fig. 2). PsychoPy (version 2023.1.3) was used to design the experimental paradigm, present the stimuli and synchronize their presentation to the EEG (Peirce et al., 2019). Stimuli were cropped to 400 x 400 pixels and were presented on a gray background of a 24.5-inch LCD screen (resolution of 1920 x 1080 pixels and a refresh rate of 120 Hz), with a viewing distance of 96 cm, secured by a chin-rest.

### Procedure

The experiment consisted of two phases (Fig. 1), which took place consecutively in a dimly lit, electrically shielded and sound-attenuated room. After placing the EEG cap on, participants read the instructions for the first, cued recall phase.

**Fig. 1.**
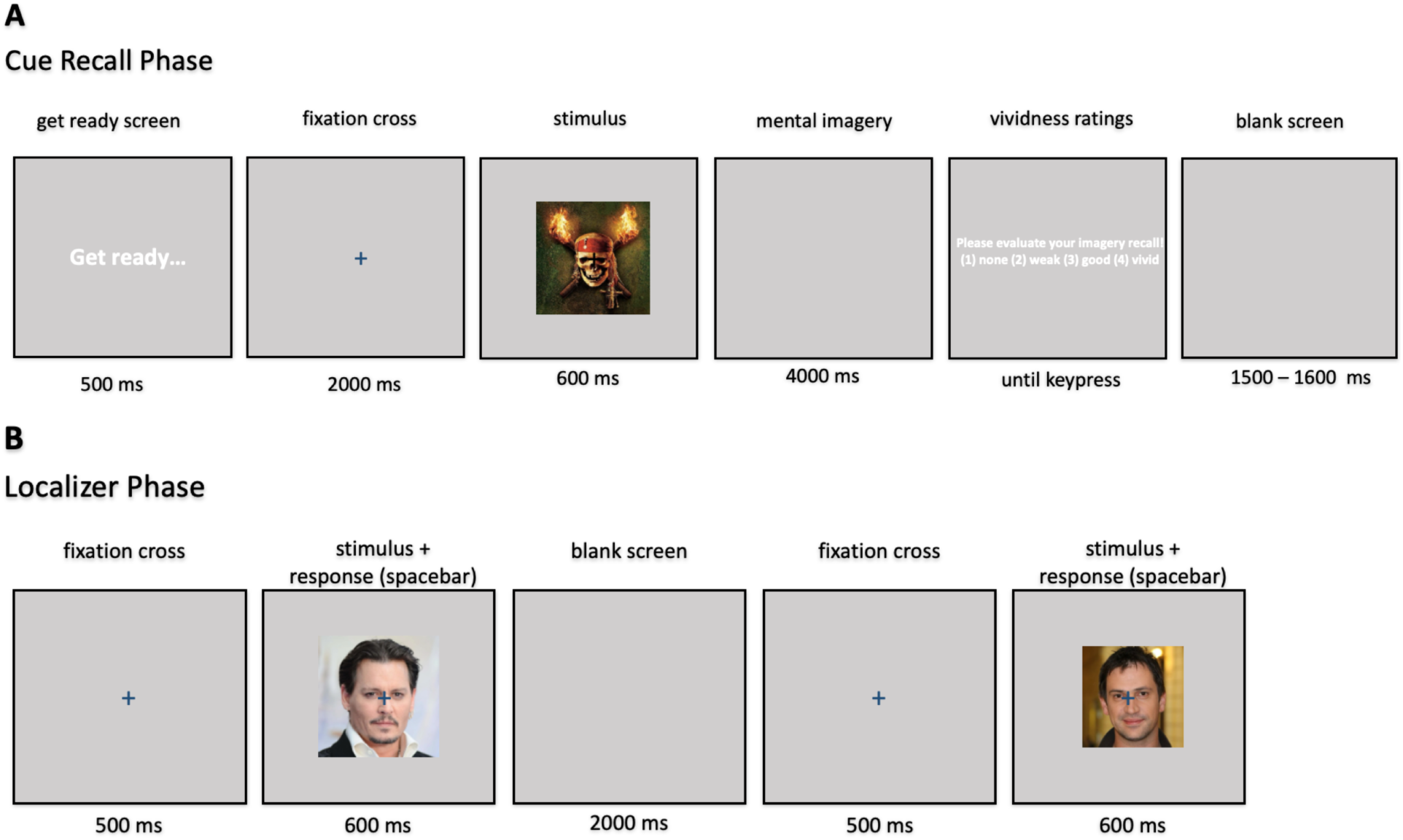
Experimental paradigm. A. In the cued recall condition, a symbol cue was presented, followed by a 4000 ms imagery period and a subsequent vividness rating. Participants were instructed to imagine and recall the face of a person, who is the most associated to this symbol. B. During the face perception “localizer” condition participants viewed faces of the persons, the most related to the symbols as well as unfamiliar control faces. This phase was executed immediately after the cued recall phase. For details see methods section.

Before each trial of, participants were presented a text to get prepared for the trial for 500 ms. Next, a fixation cross appeared in the center of the screen and remained there for the entire length of the trial. After 2000 ms, a graphic symbol was presented randomly for a duration of 600 ms, followed by a 4000 ms period in which participants were instructed to imagine the face of the most associated person to that symbol. For example, by seeing the symbol of the feature film “Iron Man”, they were instructed to recall and imagine the face of Robert Downey Jr. If the symbol was unfamiliar and did not remind them of anyone, participants were instructed not to imagine a face or person during that period. They were not instructed to imagine or maintain the visual properties of the unfamiliar cue after its offset; unfamiliar symbols served as a no-person-recall, baseline condition.

At the end of the recall period, participants had to rate the vividness and strength of their imagery recall on a Likert scale, from 1 to 4 (no imagery to vivid, strong imagery), using the number keys on a keyboard. There was no time limit for the imagery rating. The trials were ended by a blank screen and an inter-trial-interval (ITI), varied between 1500 and 1600 ms. There were five blocks of cued recall trials, each containing one repetition of the five exemplars of the eight symbols, leading to 200 trials in total. Between blocks, a short 10 s break with an on-screen countdown was presented. At the end of this phase, participants were given a form that presented each graphical symbol and were asked to name the person whom they imagined during the experiment. In the evaluation, participants reported the intended identity in 76% of the responses associated with Johnny Depp, 56% for Robert Downey Jr., 79% for Daniel Radcliffe, and 76% for Donald Trump (Supplementary Fig. 3). Because this evaluation was completed after the task rather than after every trial, it could not be used to exclude individual EEG trials.

During the second, “face localizer” phase, recorded immediately after the completion of the cued recall period, participants were presented with a fixation cross, which was followed 500 ms later by a random face image (600 ms). To make sure participants paid attention to the stimuli during the task, they were instructed to press a button when an image was presented two times in a row. Each trial was followed by a 2000 ms ITI. This localizer, face perception session was split into six blocks, each containing a repetition of the 5 exemplars of the 8 identities, resulting in 240 trials in total.

After completing the EEG sessions, participants performed a separate behavioral task to measure their familiarity with the presented identities by calculating the “mnemonic familiarity index” (MFI) (Li et al., 2022). Their subjective familiarity level with each identity was rated on a Likert scale from 0 to 9. An explicit memory score ranging from 0 to 5 was also calculated by giving one point for the correct full name, one point for occupation, up to two points for correct biographical details, and one point for a personally related episode. The mnemonic familiarity index was calculated separately for each participant and identity as:

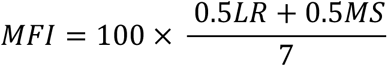

where *LR* was the subjective familiarity rating and *MS* was the explicit memory score in the same manner as Li et al. (2022).

### Behavioral analyses of the cued recall session

Trial-wise vividness ratings were averaged separately for familiar and unfamiliar symbol trials within each participant and compared by a Wilcoxon signed-rank test. A paired sample t-test was also conducted as an additional test. We also tested if vividness predicted whether participants reported the intended person. For each participant familiar identity pair, vividness ratings were averaged across the trials, and a mixed-effects logistic regression model was fitted with recall congruency as the dependent variable, centered mean vividness as a fixed-effect predictor, and random intercepts for participant and identity.

### EEG Recording and Preprocessing

The EEG was recorded by a 64-channel Biosemi Active II system with a sampling rate of 512 Hz. EEG data was preprocessed using EEGLAB (Delorme and Makeig, 2004) and custom-made MATLAB scripts. The data were band-pass filtered between 0.1 and 40 Hz, re-referenced to the average of all scalp electrodes, and downsampled to 100 Hz. Data was then epoched, based on the onset of specific image identity triggers. Epochs were extracted from -0.2 seconds prior to the trigger to 4.6 seconds post-stimulus onset in cued recall and to 1.2 seconds post-stimulus onset in the localizer sessions. Baseline correction was applied using the pre-trigger interval from -200 ms to 0 ms. No trial-level artifact rejection was performed to preserve trial numbers across the relatively long cued recall epochs and to maintain balanced classification across conditions. Previous methodological work suggests that trial rejection does not necessarily improve multivariate decoding performance when the associated data loss reduces statistical power (Delorme, 2023; Grootswagers et al., 2017; Zhang & Luck, 2025). Nevertheless, residual artifacts may have influenced decoding if they differed systematically between the experimental conditions.

### Multivariate Analysis (Decoding)

Decoding (Fig. 2) was performed using the Amsterdam Decoding and Modeling toolbox (ADAM, version: 1.14-beta; Fahrenfort et al., 2018). A k-fold cross-validation (CV) scheme was used with care taken ensuring a balanced representation of trials from each stimulus identity in the k-fold splits while keeping the number of folds at maximum. So, for the cued recall session we performed a 25-fold (200 trials / 8 different symbols = 25) CV and for the localizer session we performed a 30-fold (240 trials / 8 faces = 30) CV, for each decoding scheme. The decoded label was the binary familiarity condition (familiar vs unfamiliar) rather than the specific identity of the stimuli. Each cued recall test fold contained one trial from each of the eight symbols, corresponding to four familiar-person associated trials and four unfamiliar trials. Folds were balanced at the symbol level, but repetitions of the same visual exemplar were not explicitly grouped and could occur in different folds. Similarly, each localizer test-fold contained one trial from each of the eight identities, corresponding to four familiar and four unfamiliar face trials. We trained a linear discriminant analysis (LDA) classifier for each time-sample separately on all folds except one and tested the decoder on the held-out fold in an iterative fashion so that every fold was used once for testing. The decoding accuracies were then averaged across all folds and across participants, separately for each timepoint, to lead to the final decoding accuracy as the function of time.

**Fig. 2.**
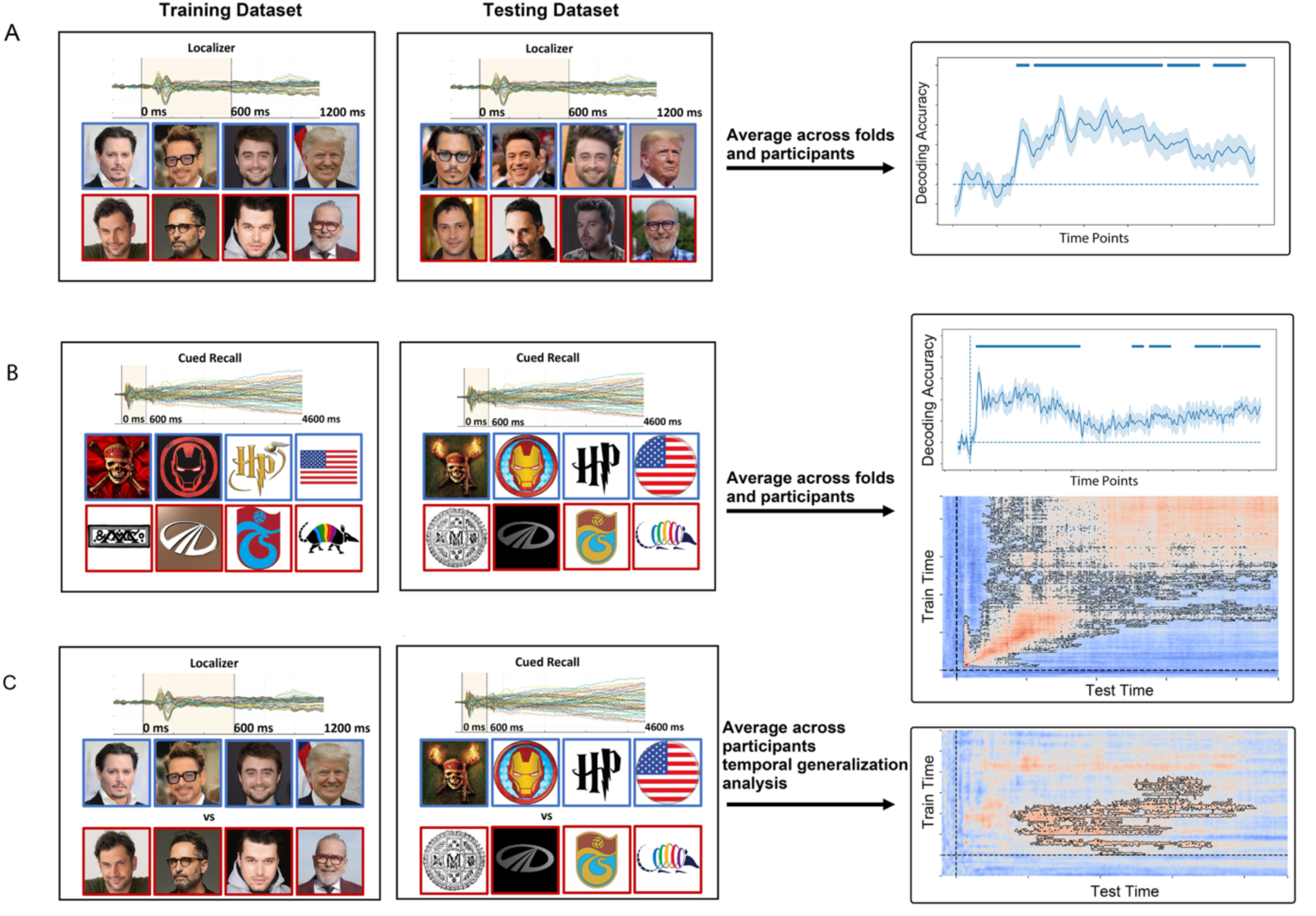
The logic of the decoding analyses. **(A)** Perceptual face “Localizer” trials: time-resolved decoding. Sensor-level EEG from the presented faces (0-1200 ms post-stimulus) was used to train and test a classifier, distinguishing familiar (blue frames) vs unfamiliar (red frames) faces. A 30-fold, within-participant cross-validation scheme was applied. The right side shows plots of the group-average accuracies over time; dotted red line marks chance level **(B)** Cued recall/imagery trials: time-resolved decoding and temporal generalization. We performed familiar vs unfamiliar decoding using 25-fold, within-participant cross-validation (time course shown), and then computed the temporal-generalization matrix by training at each time point and testing at every time point within the cued recall data. **(C**) Cross-domain generalization of familiarity (localizer -> cued recall/imagery). A cross-domain temporal-generalization analysis assessed whether familiarity representations, learned during face perception in the localizer trials generalize to the symbol-to-face imagery trials. Classifiers were trained at each localizer time point and tested at each cued recall time point; the resulting train x test matrix was averaged across participants. Warmer colors indicate higher-than-chance generalization. Red outlines indicate significant clusters. Stimulus frames show examples of the stimuli; blue/red frames denote familiar/unfamiliar exemplars. All accuracies reflect sensor-level multivariate patterns, averaged across folds and then participants. Statistical evaluations were done by 10000 cluster-based permutations across time and train x test time.

Decoding was performed both on the data of the entire scalp as well as in six separate, pre-defined topographic regions of interest (ROI), including the right and left frontal, central-temporal and parietal-occipital cortex (Ambrus et al., 2019). The left anterior ROI included Fp1, Fpz, AF3, AFz, F7, F5, F3, F1, Fz, FC3, FC1, and FCz; the right anterior ROI included Fp2, Fpz, AF4, AFz, F8, F6, F4, F2, Fz, FC4, FC2, and FCz. The left central ROI included FT9, FT7, TP9, TP7, C5, C3, C1, Cz, CP3, CP1, CPz, and T7; the right central ROI included FT10, FT8, TP10, TP8, C6, C4, C2, Cz, CP4, CP2, CPz, and T8. The left posterior ROI included PO9, PO7, PO3, POz, P9, P7, P3, P1, Pz, O1, Oz, and Iz; the right posterior ROI included PO10, PO8, PO4, POz, P10, P8, P4, P2, Pz, O2, Oz, and Iz. Midline electrodes were included in both corresponding left and right ROI groups. A schematic of the six ROIs is available in Supplementary Fig. 4.

Familiarity decoding was performed in both phases of the paradigm: during cued recall the classifier discriminated familiar person-associated symbol trials from unfamiliar/no-person-recall symbol trials, while in the localizer phase it discriminated familiar versus unfamiliar faces. We performed these analyses both in a time-resolved fashion (training on each specific timepoint of trials and testing on the same timepoints, resulting in a one dimensional, diagonal vector of classification accuracies, consisting of every timepoint in the trial) as well as in a temporal cross-generalization manner (training the decoder on each timepoint and testing it on every other timepoint, resulting in a 2D training time x testing time temporal generalization matrix of classification accuracies). Additionally, we performed cross-condition decoding analyses by training the classifier on localizer data and testing on retrieval data in a temporal cross-generalization manner. There was no cross-validation on the cross-condition decoding analyses, as the training and testing datasets were independent from each other. The perception-to-recall training-test direction was used as the primary cross-domain analysis because the localizer session provided a perceptually well-defined training dataset in which both familiar and unfamiliar faces were physically presented. This allowed us to test whether familiarity-related patterns, learned during face perception generalize to later processing during cued recall or not. A complementary reverse direction analysis, in which classifiers were trained on the cued recall data and tested on the face perception data was also conducted merely as a robustness check (Supplementary Materials).

### Statistical testing

We calculated group level decoding accuracies from individual first-level results. For the time-resolved decoding, performance was compared against chance level (50% for familiarity decoding), for each time point separately, using one-tailed one-sample t-tests (decoding accuracy should not be below chance, see Kaiser et al., 2016; Bae, 2020) across participants. To deal with multiple comparisons, a non-parametric, cluster-based permutation test was applied, across time points with 10000 iterations (Maris & Oostenveld, 2007). This method used an uncorrected significance threshold of *p* < .05 to identify potential cluster members first, followed by a corrected cluster-level threshold of *p* < .05 to measure the overall significance of the identified clusters.

For temporal generalization analyses, the cluster-based permutation approach was extended to the 2D matrix to identify significant (including off-diagonal) clusters, using the same significance thresholds.

For visualization purposes, the resulting group average decoding accuracy time courses were smoothed, using a 3 timepoint (30 ms) moving average. Statistical analysis was performed separately for the decoding results of the entire scalp and the six pre-defined ROIs.

## Results

### Localizer paradigm: familiarity decoding for faces

To confirm that our participants were indeed familiar with the selected famous persons, we first estimated the behavioral MFI for each identity separately. Fig. 3 shows that the participants showed significantly higher familiarity scores for the four familiar persons, compared to the unfamiliar controls (paired *t*-test: *t* = 37.5, *p* < .00001, Cohen’s *d* = 2.75). Second, we performed a binary familiar versus unfamiliar classification on the EEG data, during the localizer phase of the experiment, over the entire scalp as well as in the six pre-defined ROIs separately (Fig. 4). Accuracies for familiarity decoding were significantly above chance for the entire scalp in four temporal clusters, with the first between 100 and 160 ms (peak at 110 ms, *p* < 0.05, Cohen’s *d* = 0.64), the second between 180 and 770 ms (peak at 300 ms, *p* < 0.0001, Cohen’s d = 0.82), the third between 790 and 940 ms (peak at 860 ms, *p* < 0.01, Cohen’s *d* = 0.84), and the last between 1000 and 1150 ms (peak at 1010 ms, *p* < 0.01, Cohen’s *d* = 0.53).

**Fig. 3.**
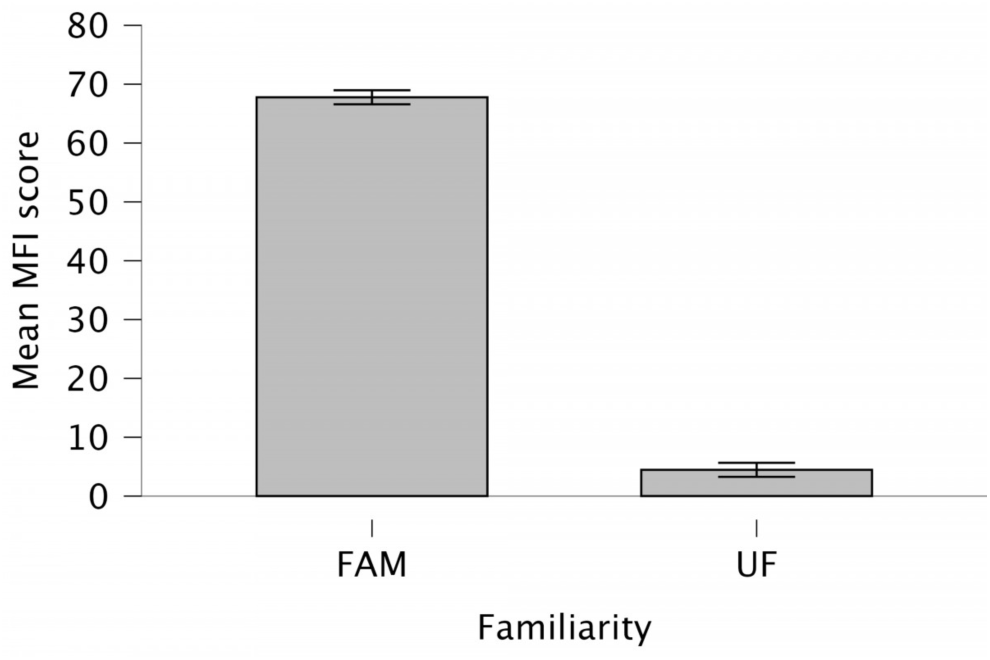
Mean face familiarity index (MFI) scores for the familiar (FAM) and unfamiliar (UF) identities. Bars represent the average MFI values, computed across participants and stimulus identities. Error bars indicate the standard error of the mean (SEM).

**Fig 4.**
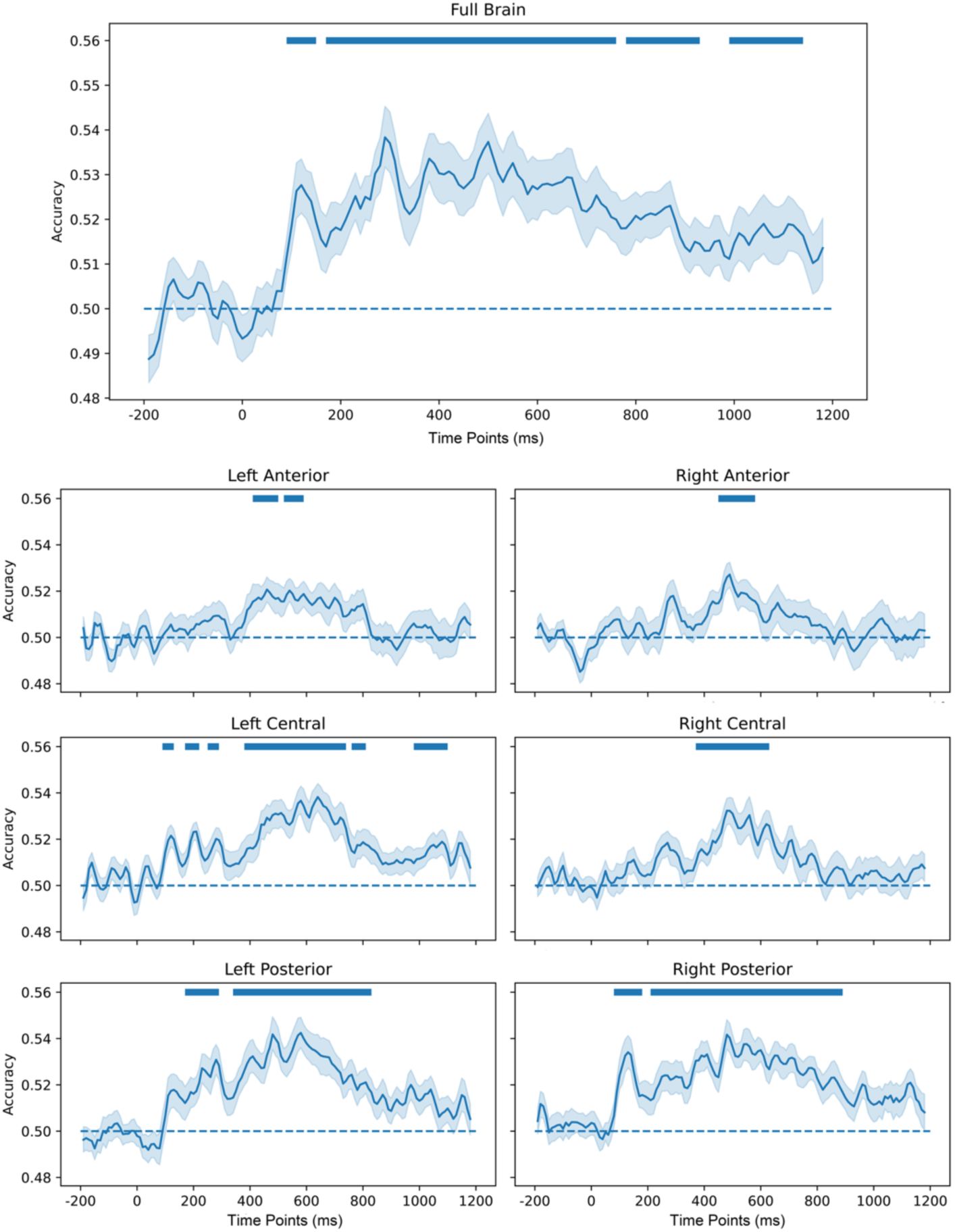
Group-level time-resolved familiarity decoding accuracies from the face perception trials. Horizontal lines indicate the clusters of timepoints with significantly higher decoding accuracies when compared to the chance level. Decoding accuracy is plotted as a proportion; 0.50 corresponds to 50% accuracy and to chance level performance. Significance was tested by 10000 one-tailed cluster-based permutations, p< 0.05.

Similarly, when estimating familiarity information within the pre-defined ROIs, the temporal dynamics of the decoding results were in line with previous findings on familiarity representation (Fig. 4). Familiarity could be decoded in bilateral posterior ROIs the earliest (from 180 ms over left and from 90 ms over right hemisphere) and remained (with short interruptions) significant in a prolonged period until 840 ms and 900 ms post-stimulus onset over the left and right hemisphere, respectively (for detailed statistics see Table 1). It was shorter and had a later onset over central and anterior ROIs (except for left central electrodes where it started at 100 ms and went until 1110 ms with disruptions; Table 1). These temporal dynamics are in correspondence with prior MVPA and ERP results, regarding familiarity for experimentally familiarized (Ambrus et al., 2021; Andrews et al., 2017; Dalski et al., 2022, 2023), famous (Ambrus et al., 2019, Dobs et al., 2019) as well as for personally familiar faces (Ambrus et al., 2021 and Li et al., 2022; Wiese et al., 2024). Overall, this suggests that our participants were indeed familiar with the faces of the selected famous persons.

**Table 1.** Results of the time-resolved familiarity classification for faces.

| ROI | Onset (ms) | Peak (ms) | End (ms) | Cluster-based $p$-value | Cohen's $d$ (peaks) |
| --- | --- | --- | --- | --- | --- |
| Full Brain | 100 | 110; 300; 860; 1010 | 1150 | 0.023; <0.0001; 0.005; 0.009 | 0.64; 0.82; 0.84; 0.53 |
| Left Anterior | 420 | 470; 540 | 600 | 0.016; 0.02; | 0.65; 0.63 |
| Right Anterior | 460 | 490 | 590 | 0.007 | 0.73 |
| Left Central | 100 | 120; 200; 290; 580; 790; 1080; | 1110 | 0.037; 0.024; 0.046; <0.001; 0.041; 0.009 | 0.60; 0.90; 0.80; 0.86; 0.54; 0.55 |
| Right Central | 380 | 490 | 640 | <0.001 | 0.86 |
| Left Posterior | 180 | 280; 570 | 840 | 0.009; <0.0001 | 0.82; 0.95 |
| Right Posterior | 90 | 130; 480 | 900 | 0.012; <0.0001 | 0.84; 1.04 |

### Behavioral results of the cued recall session

Imagery vividness was significantly higher for familiar (M = 3.16, SEM = 0.10) than for unfamiliar symbol trials (M = 1.80, SEM = 0.14), (Wilcoxon signed-rank test (*W* = 553, *p* < .001; Supplementary Fig. 5A)). A complementary paired sample t-test led to the same conclusion (*t*(33) = 9.41, *p* < .001, Cohen’s *d* = 1.61). A mixed-effects logistic regression additionally showed that vividness significantly predicted whether participants reported imagining the intended person or not (β= 1.86, SE = 0.40, *p* < .001; Supplementary Fig. 5B). These findings indicate that the familiar and unfamiliar conditions differed in imagery/person recall engagement.

### Familiarity decoding for abstract symbols

One of our major aims was to test if the neural representations of familiarity for abstract, graphical symbols can be decoded from the EEG signal, similarly to faces. To test this, for each participant and time point, we trained an LDA classifier to discriminate between the patterns of neural signals evoked by familiar and unfamiliar symbols, presented as cues, during the cued recall trials (Fig. 5).

**Fig 5.**
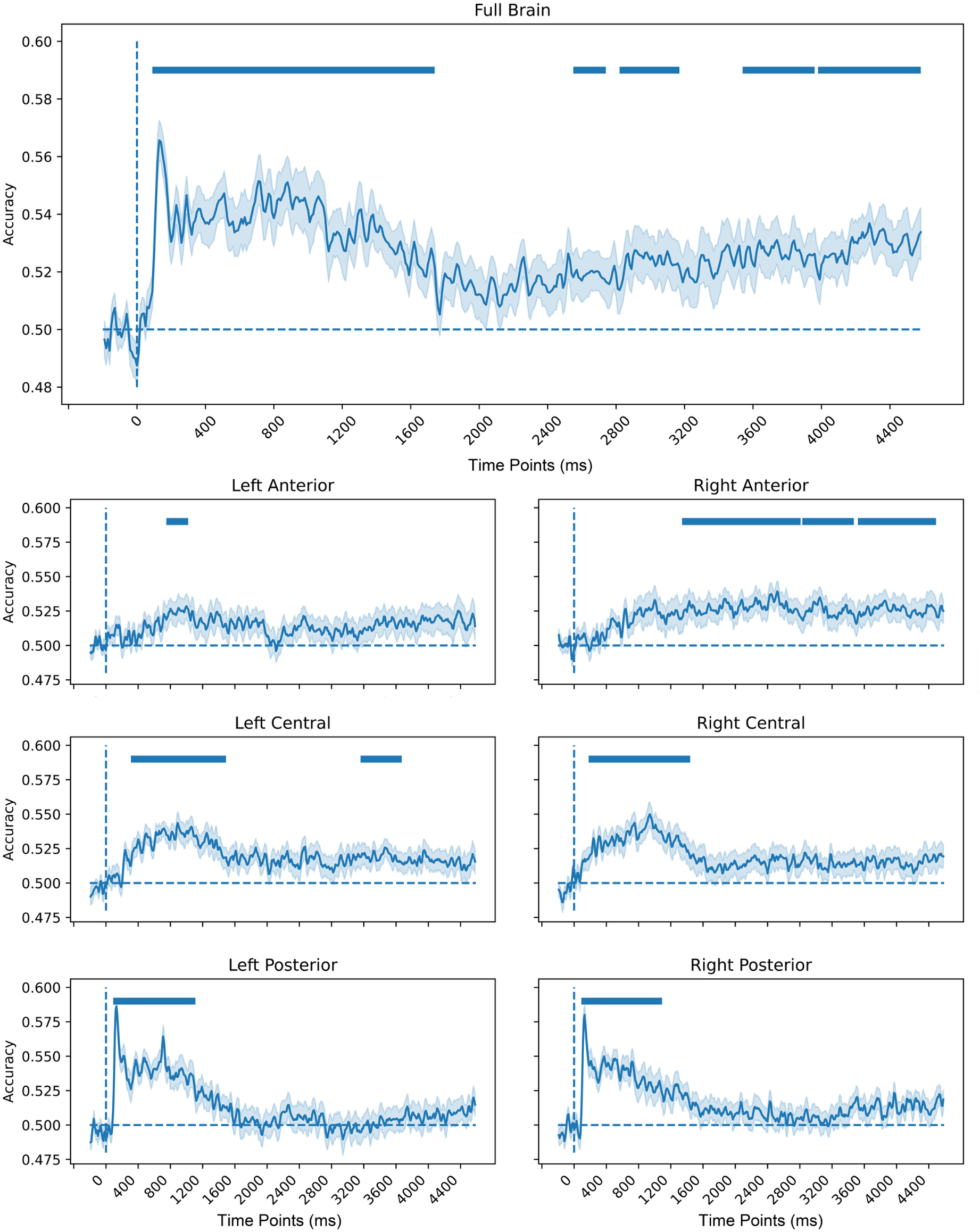
Group-level time-resolved familiarity decoding accuracies from the cued recall trials. Horizontal lines below the figures indicate timepoints with significantly higher decoding accuracy when compared to chance level (dotted line). Decoding accuracy is plotted as a proportion; 0.50 corresponds to 50% accuracy and to chance level performance. Significance was tested by 10000 one-tailed cluster-based permutations, p< 0.05.

Decoding accuracies were significant in five clusters between 100 and 1750 ms (peak at 130 ms, *p* < 0.0001 and Cohen’s *d* = 1.58), 2560 and 2750 ms (peak at 2690 ms, *p* <0.05 and Cohen’s *d* = 0.58), 2830 and 3180 ms (peak at 2910 ms, *p* < 0.05 and Cohen’s *d* = 0.67), 3550 and 3970 ms (peak at 3830 ms, *p* < 0.05 and Cohen’s d = 0.59), and 3990 and 4590 ms (peak at 4270 ms, *p* < 0.01 and Cohen’s *d* = 0.88), when tested across the entire scalp (Fig. 5).

Decoding of symbol familiarity in the pre-defined ROIs revealed significant familiarity information in almost every electrode cluster, but with temporal dynamics that differed from those observed for faces (Fig. 5). Familiarity information was present relatively early in the bilateral posterior ROIs, starting at 100 ms in both hemispheres, and remained significant until 1120 ms and 1100 ms over the left and right posterior ROIs, respectively (Table 2). The central ROIs showed significant familiarity information starting later (at 320 and 190 ms for the left and right hemisphere, see Table 2) and ended at 1500 ms and at 1450 ms on the left and right hemispheres (with an additional cluster between 3170 and 3680 ms on the left side). The left anterior ROI showed significant decoding only for a short time between 760 and 1030 ms. Finally, the right anterior ROI also showed significant familiarity information very late (1350 to 4500 ms; with gaps between 2820-2840 ms, and between 3480-3530 ms; see Table 2).

**Table 2.**
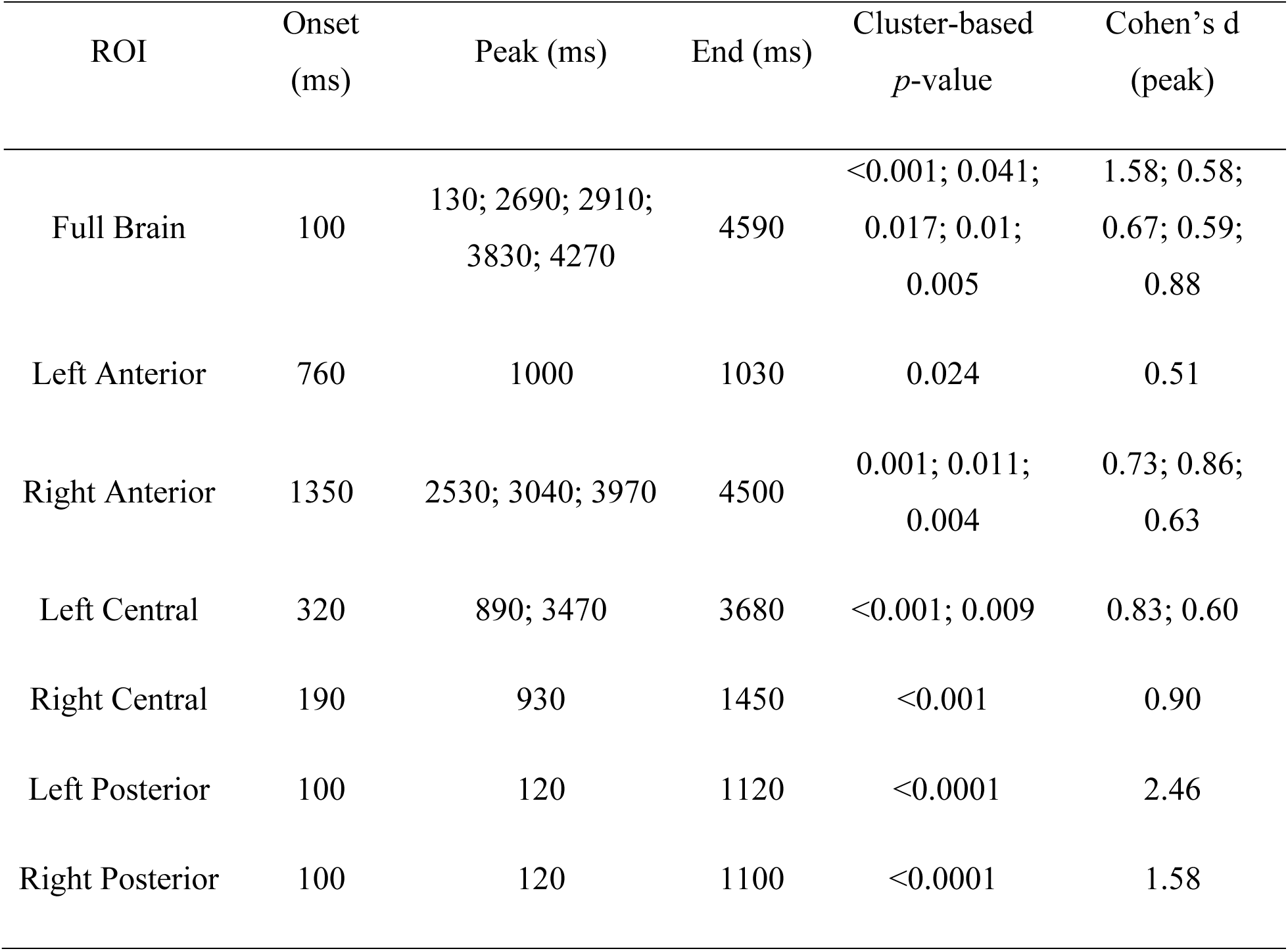
Results of the time-resolved familiarity classification for symbols.

Interestingly, while the posterior and central ROIs showed discrete peaks of classifier performances, just like various previous studies for faces (Ambrus et al., 2019; Dobs et al., 2019), the right lateralized anterior ROI showed a sustained, tonic decoding performance, extending to several seconds. This activation had no temporal overlap with that of posterior electrode clusters and only a short overlap with central electrode clusters (150 and 100 ms with left and right). This result is in line with the findings of a recent study, showing similarly late and prolonged category representations during the cued recall of people vs. places (Corriveau et al., 2023) and adds the information that it originates dominantly from the anterior part of the right hemisphere. Together, these results suggest that the activation of a person representation from abstract symbols is delayed and reflects different neural processing when compared to sensory stimuli.

To test the stability of the temporal dynamics of familiarity representations, elicited by the abstract symbols, we conducted a temporal-generalization analysis, using the same group-level decoding procedure as the previous analysis. For this, we trained the classifiers on one time point and tested on all other time points (King and Dehaene, 2014). Figure 6 presents the temporal generalization matrices for the entire scalp as well as for the six pre-defined ROIs, separately. The data of the entire scalp shows several distinct stages of familiarity representation. During the first prolonged stage (90 to around 1800 ms), classifiers showed significant decoding along the diagonal, with relatively brief off-diagonal periods, suggesting a rapidly evolving, but time-specific familiarity representation. From around 2000 ms onwards until the end of the trial there was a broad off-diagonal generalization pattern, potentially reflecting the stabilization of the familiarity representation of recalled items. The temporal generalization of the separate ROIs revealed the differential involvement of the two hemispheres in this representation. While the right anterior ROI showed a pattern that started relatively late at around 1500 ms and showed a very broad temporal generalization until the end of the trial, for the central ROIs we observed an earlier start (around 200 ms) that broadly follows the diagonal with a temporal generalization taking only place after 1400 ms. The significant cluster on left posterior ROI on the other hand, was the earliest (90 ms to approximately 2000 ms), following a diagonal pattern. Overall, temporal generalization reveals a transition from early dynamic to later, sustained familiarity encoding. Early, time-specific signals (e.g. shown by the strong diagonal pattern at the start in the posterior and central ROIs) give way to a later, tonic, right anterior cluster localized state, that supports maintaining and acting on the retrieved item.

**Fig 6.**
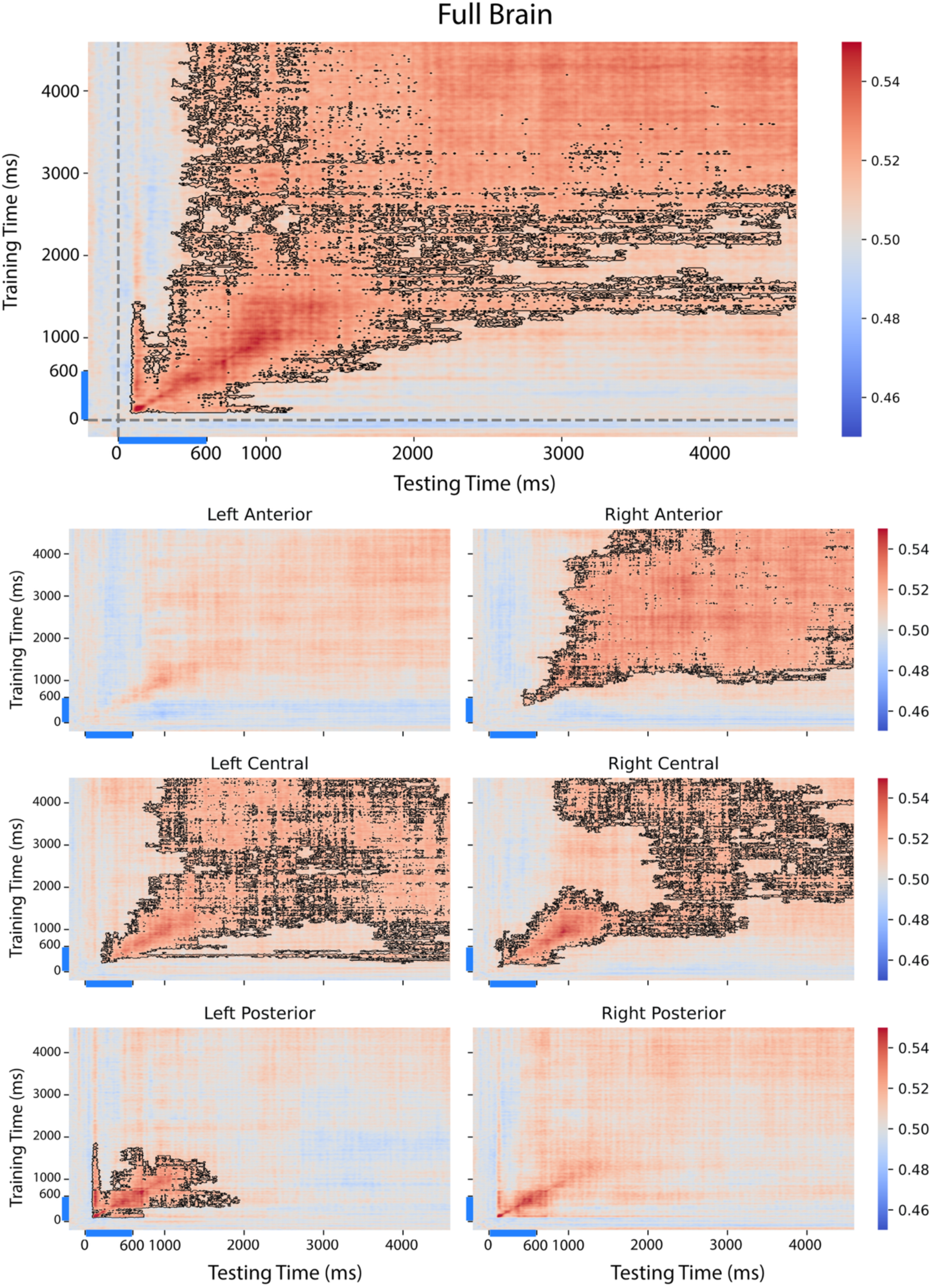
Temporal generalization of familiarity information for the cued recall trials. Horizontal rows indicate timepoints the classifiers were trained on and the vertical columns indicate timepoints the classifier was tested on. Decoding accuracy is plotted as a proportion; 0.50 corresponds to 50% accuracy and to chance level performance. Significant temporal clusters are indicated with the surrounding black lines. Significance was tested by 10000 one-tailed cluster-based permutations, p< 0.05.

### The temporal overlap of familiarity representations during perception and imagery

To investigate whether perceptual familiarity processing generalizes to and overlaps with that of imagery, we ran a cross-domain familiarity decoding analysis. A classifier was trained on the faces of the perceptual phase trials and tested on the data of cued recall trials (Fig. 7). As the length of the trials was different for these two conditions, we followed the methods of Dijkstra and colleagues (2018) and only tested temporal generalization.

**Fig 7.**
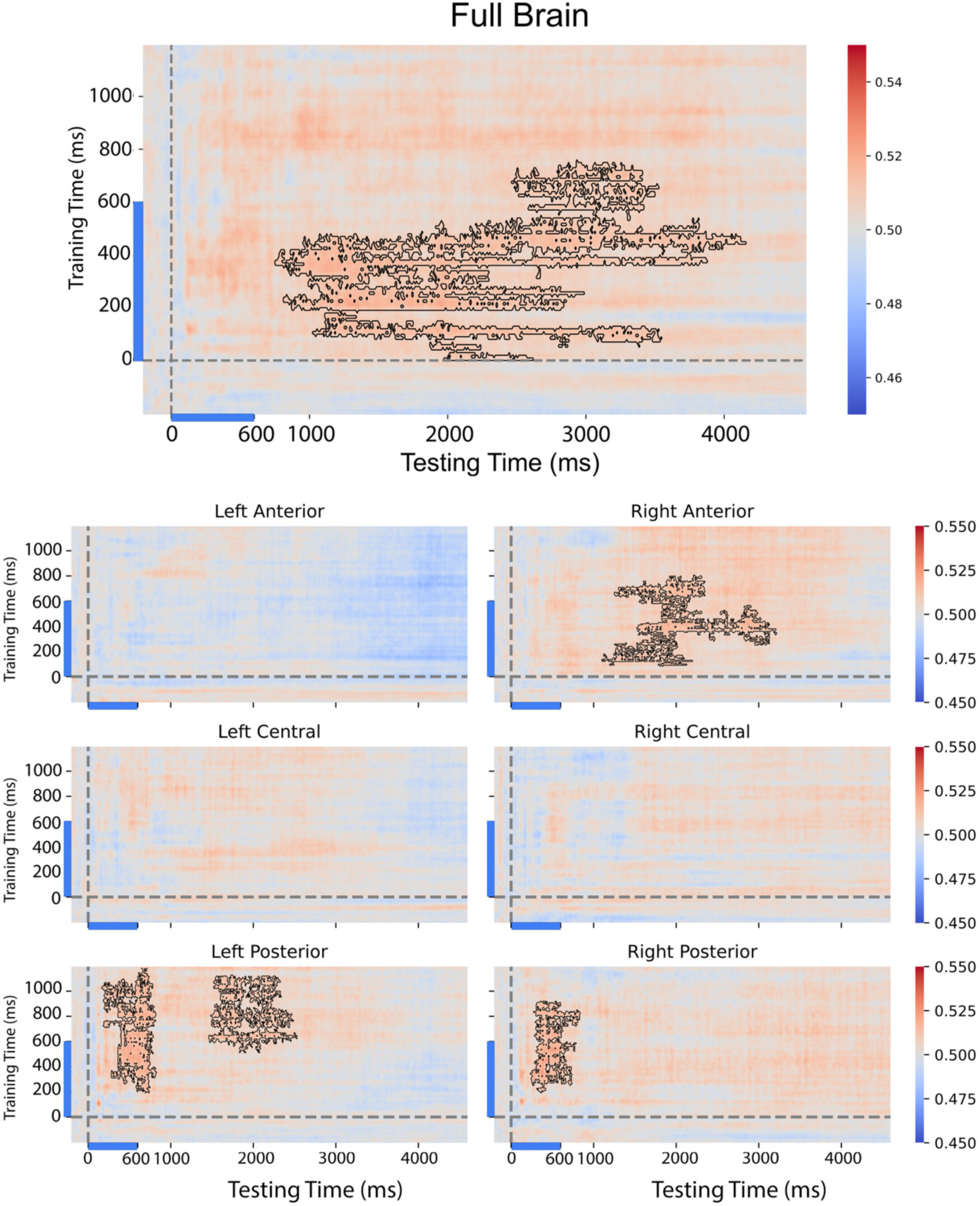
Temporal generalization of familiarity between perception and symbol-cued recall/imagery related processing. Horizontal rows indicate timepoints that the classifiers were trained on (perceptual data of the localizer trials) and vertical columns indicate timepoints when the classifier was tested (data from the cued recall trials). Decoding accuracy is plotted as a proportion; 0.50 corresponds to 50% accuracy and to chance level performance. Significant temporal clusters are indicated with the surrounding black lines. Significance was tested by 10000 one-tailed cluster-based permutations, p< 0.05.

There were significant clusters in the whole brain temporal generalization matrix as well in the right anterior and bilateral posterior ROIs. Considering the entire scalp, we found that perceptual information between 10 and 760 ms generalized to imagery in a long time-window between 760 and 4160 ms. This cluster mostly corresponded to the face stimulus presentation period in the localizer condition and the post-cue recall/imagery period in the cued recall trials. In addition, the posterior ROIs showed generalization from the perceptual processing between 200 - 1190/920 (left/right hemispheres) to symbolic cue-evoked activity between 170 - 830 ms on the left and 240 - 840 ms on the right. This suggests the strong temporal overlap of face and graphic symbol evoked familiarity representations and support further prior findings regarding the category-independent nature of familiarity representation (Klink et al., 2023). Additionally, the left posterior ROI showed a second cluster, trained on perceptual data between 520 and 1130 ms, generalized to imagery data between 1470 - 2530 ms. Importantly, the right anterior ROI showed entirely different temporal dynamics of generalization when compared to the posterior electrode clusters. Perceptual familiarity processing between 90-810 ms generalized to symbol-evoked processing between 1110 and 3220 ms. This generalization period corresponds to what we found for the same ROI when analyzing the cued recall trials (see Fig. 5).

To rule out the possibility that the results were driven by participants who frequently imagined people in response to unfamiliar symbols (contrary to the instructions), we repeated the cross-domain decoding analysis after excluding seven participants (*N* = 27) who reported imagining a person for more than two unfamiliar symbols. Nonetheless, the main pattern of results (Supplementary Fig. 6) remained comparable to the original analysis. The full results of this additional analysis are reported in the Supplementary Materials.

The complementary reverse train-test direction (classifier trained on cued recall and tested on perceptual localizer face data) analysis revealed broadly convergent cross-domain generalization in the bilateral posterior and right anterior ROIs (Supplementary Fig. 7).

Finally, sensitivity analyses using less conservative statistical thresholds reproduced the primary cross-domain temporal generalization pattern. Lowering the cluster-level threshold to *p*<0.1 did not change the pattern of significant results, whereas using a more liberal pointwise cluster-forming threshold of *p*<0.1 produced the same overall spatiotemporal pattern, with some clusters increasing in size, although the right posterior effect no longer survived cluster-level correction (Supplementary Fig. 8).

## Discussion

The major results of the current article are the following: (1) Familiarity decoding is possible for abstract symbols with similar spatio-temporal dynamics to those for faces. This processing has two separate, spatio-temporally non-overlapping stages: while the first ∼1500 ms is relatively time-specific, the later periods, located over right anterior sites generalize strongly across time. (2) Perceptual familiarity processing generalizes to symbol-cued recall/imagery related processing. The two separate processing stages are also present in their generalization. While early, bilateral posterior face familiarity processing generalizes to cue-evoked activity in a temporally overlapping manner, the later, right frontal (and left posterior) localized perceptual familiarity generalizes only to much later imagery data in a temporally non-overlapping manner.

Overall, our results suggest the transition of familiarity processing from an early, dynamic processing, distributed over posterior scalp sites (potentially corresponding to occipito-temporal and parietal areas, reflecting perceptual processing) towards a later, sustained familiarity encoding over the right anterior scalp sites (potentially corresponding to frontal areas), that is compatible with successful recall and imagery of concepts.

### Similar temporal dynamics of familiarity processing for faces and abstract symbols

During cued recall trials, neural activity likely reflects both perceptual analysis of the cue and imagery of the associated face. Significant information was found during and immediately after the presentation of the stimuli across the scalp, except for the right anterior electrode cluster, where it occurred much later, from 1100 ms onwards.

Distinguishing perceptual and imagery-related components is challenging in this type of paradigm. Importantly, the familiar and unfamiliar cued recall conditions differed not only in familiarity but also in the expected engagement of person retrieval and imagery. The present contrast may therefore reflect a combination of symbol familiarity, person-related semantic retrieval, imagery engagement, and retrieval monitoring. Accordingly, these effects can also be interpreted as familiarity-related processing during symbol-cued person recall rather than as a pure measure of familiar face imagery.

Because three of the familiar symbols were related to famous movies, the symbols may also have elicited imagery of scenes, places, or contextual events from the movies in addition to the instructed person imagery. We did not collect reports of scene or contextual imagery and cannot exclude their contribution either. Consequently, the cued recall effects may additionally include person imagery, semantic/contextual associations, and scene-related imagery (Scrivener et al., 2025).

Nonetheless, we argue that while the early, transient and time-specific decoding dominantly reflects perceptual processing, the later, tonic and temporally widely generalizing information is compatible with imagery related processing of the recalled objects. First, the temporal dynamics of familiarity information reflect those of many previous uni and multivariate studies (for a review see Wiese et al., 2024). Familiarity information prior to 2000 ms peaked at around 130 ms as well between 400-1000 ms. This is similar, albeit somewhat later, when compared to faces and scenes (Klink et al., 2023). The spatial distribution of this early familiarity encoding was focused on bilateral posterior parts of the brain, again supporting previous face-related familiarity findings. This posterior scalp distribution is also broadly consistent with fMRI evidence showing medial parietal involvement during recall of familiar people and places and a posterior network linking scene perception with memory for familiar places (Silson et al., 2019a; Steel et al., 2021). However, the present sensor-level EEG results do not allow us to specifically attribute the effects to these cortical regions. In addition, the temporal generalization analysis showed that this effect is restricted to the diagonal, suggesting a very time-specific processing. All these findings are compatible with the interpretation of feed-forward, perceptual processing of familiarity in this time-window.

Unlike the early familiarity representation, the information present after 1350 ms post-stimulus onset was prolonged, tonic and localized to the right anterior electrode cluster, without almost any spatio-temporal overlap with the former one. This information generalized across time very broadly, suggesting robust, but less time-specific processing. These results are compatible with those found for stimulus category representations. Dijkstra et al., (2018) found that face-scene category imagery related information generalized broadly over time. However, category imagery information had a far earlier onset (around 500 ms), when compared to the familiarity imagery onset of the current study. In addition, they found the effect localized dominantly to the posterior parts of the brain, with very little frontal components. This conclusion was also supported by Xie et al (2020) who found shared perceptual and imagery information about object categories in the alpha frequency band, restricted to posterior electrode sites. As imagery activates a large network of areas (Ganis et al., 2004, Ishai, 2000) we argue that this spatio-temporal difference is due to the different location of category and familiarity recall and imagery in the brain. This suggests that imagery processing depends on its content and on the task at hand, a theory worth to test explicitly within the same subjects in the future.

Regarding the origin of the later, frontal familiarity imagery information there are at least two alternative explanations. First, it is possible that the imagined faces are indeed represented in the right frontal regions. This is, however, at odds with some prior studies which found that imagined and perceived stimulus categories are represented similarly (Xie et al., 2020). To the best of our knowledge this is the first study testing imagery, related to familiarity representation and differences between stimulus category and familiarity imagery may explain the different spatial distribution of the effects. Second, our findings can be interpreted in the frame of the “right frontal old/new effect”, as well; retrieval monitoring and decision-making processes have been associated with the right frontal old/new effect (Wilding & Rugg, 1996; Hayama et al., 2008), which is compatible with the late and sustained time course of the present right anterior ROI pattern. A broader contribution of the right hemisphere to familiar person recognition may also play a part in this asymmetry (Gainotti, 2007). Because traditional sensor-level EEG recording does not allow us to have precise cortical localization, we interpret this pattern as potentially reflecting a combination of imagery maintenance, higher-level person related processing, and retrieval evaluation.

### Familiarity processing generalization from perception to imagery

First, we found a significant cross-domain decoding performance across faces and symbols around stimulus presentation and before 800 ms post-stimulus onset over bilateral posterior areas. This supports further the similar neural processing when viewing faces or the symbolic cues of the imagery trials. This finding is also consistent with recent EEG and fMRI results and suggest that familiarity is encoded in an abstract form, generalizing across stimulus categories (Klink et al., 2023; Castro-Laguardia et al., 2024). For example, Klink et al (2023) showed temporal generalization across faces and scenes at a similar onset (around 200 ms). Similarly, Castro-Laguardia et al. (2024) identified abstract familiarity regions in which patterns discriminating familiar from unfamiliar faces could also discriminate familiarity for names. Taken together, these studies support category and modality independent processing of familiarity-related information and extends it toward imagery recall.

We also observed a much later, sustained cross-decoding temporal generalization cluster, localized to the right anterior areas, when classifiers were trained on perceptual face data and tested on the cued recall data. This generalization was present already from 90 ms post face stimulus onset and suggests also a representational overlap between perception and recall/imagery-related processing. It fits well with accounts that emphasize post-retrieval monitoring/decision and maintenance operations during recall, as we also mentioned above (Hayama et al., 2008; Wilding & Rugg, 1996). This finding is also in line with those of Dijkstra et al (2018) who suggested that imagery is a top-down process that bypasses early sensory stages of perception and enters more anterior, higher-level perceptual processing stages directly. The presence of this right anterior temporal generalization cluster in our data is at accord with this interpretation and localizes the processing to the right hemisphere. The timing and the prolonged duration of this cluster are also in line with prior work that investigated cued recall of familiar persons and scenes (Corriveau et al., 2023). In addition, research directly contrasting self-generated and cue-induced imagery indicates that both modes engage anterior and posterior cortices; however, cue-induced imagery tends to show relatively stronger posterior involvement, whereas self-generated imagery shows relatively stronger anterior engagements (Hu & Yu, 2023). Important to note, however that their results were obtained with low-level stimuli which may have amplified posterior contributions.

Related fMRI work has shown a posterior-anterior distinction within medial parietal cortex, with posterior regions responding more strongly during scene perception and relatively more anterior regions responding more strongly during memory-based scene construction (Silson et al., 2019b). Together with evidence for distinct medial parietal subdivisions during familiar-person and familiar-place recall (Silson et al., 2019a) and for place-memory regions positioned anterior to scene-perception areas (Steel et al., 2021), these findings support a general distinction between visually driven and memory-related processing stages.

However, the above interpretation does not exclude the possibility that the right anterior activity also reflects a shared, higher-order representational format during imagery. Previous studies showed substantial perception-imagery overlap in representational geometry (Dijkstra et al., 2018; Xie et al., 2020), and that category specific familiarity information can be decoded during imagery (Ragni et al., 2021). Overall, the late, right anterior familiarity generalization effect may be comprised of a mix of control/monitoring/executive and higher-level representational processes that work together when a person engages in imagery tasks (Corriveau et al., 2023; Dijkstra et al., 2018, 2019).

A second, later left posterior temporal generalization also emerged from our cross-domain decoding analysis. Classifiers trained on face perception during the period after observing faces (520-1130) ms generalized to imagery at 1470-2530 ms. This can be considered as a form of delayed reinstatement of perceptual familiarity codes in ventral occipito-temporal cortex which would be supported by earlier research showing that imagery activates category specific high level visual areas such as the fusiform face area as well (Ishai et al., 2000, 2002; O’Craven & Kanwisher, 2000). Similarly, accounts of a left fusiform imagery node that bridges visual and semantic information during visual mental imagery (Spagna et al., 2021, 2024) and neuropsychological evidence implicating this region and its connectivity in aphantasia (Kutsche et al., 2025; Liu et al., 2025) are also compatible with our data considering that the symbolic cues in our paradigm may have led to imagery of faces through semantic person identity knowledge or naming the persons. Future studies can utilize effective connectivity methods and analyses in the frequency domain to better characterize the information flow during imagery and perception. They can also control for aspects such as imagery vividness or can find ways of interfering with executive/control processes during the task to probe the exact role of right anterior areas.

We did not collect independent valence or arousal ratings for the identities or symbols. Therefore, affective, semantic, and sociopolitical differences between the familiar and unfamiliar stimuli may have contributed to the measured neural differences. We also did not administer a standardized imagery questionnaire and therefore cannot determine if the sample included participants with aphantasia or unusually weak imagery. The trial-wise vividness ratings provide a task specific measure but cannot be used to identify or rule out aphantasia.

## Conclusion

Our study extends the results of earlier work on cued recall and familiarity representation by showing that familiarity representations elicited by symbolic cues evolve from dynamic category-independent posterior activity into sustained right frontal processing, bridging perceptual and imagery-related recall representations. Overall, our findings point to familiarity-related information processing to be both robust across different types of stimuli and flexible across perception and imagery.

## CRediT authorship contribution statement

**Yeliz Dinç:** Conceptualization, Methodology, Software, Validation, Formal analysis, Investigation, Data curation, Writing – original draft, Writing – review & editing, Visualization, Project administration. **Yang Shi:** Software, Visualization. **Gyula Kovács:** Conceptualization, Methodology, Software, Resources, Writing – original draft, Writing – review & editing, Visualization, Supervision, Project administration, Funding acquisition.

## Declaration of competing interest

The authors declare that they have no competing interests.

## Data and materials availability

Data and all materials are available on Figshare (https://figshare.com/projects/The_Neural_Dynamics_Of_Familiarity_Representation_During_Cued_Recall/280074). The study and analysis design were not preregistered.

## Acknowledgments

We thank Sophia Gommlich and Azlia Istiqomah for their support with participant recruitment and EEG data acquisition. We also thank Bettina Kamchen for her expert technical assistance during setup and recordings.

## Notes

https://figshare.com/projects/The_Neural_Dynamics_Of_Familiarity_Representation_During_Cued_Recall/280074

### Competing Interest Statement

The authors have declared no competing interest.

## References

Ambrus, G. G., Eick, C. M., Kaiser, D., & Kovács, G. (2021). Getting to Know You: Emerging Neural Representations during Face Familiarization. The Journal of Neuroscience, 41(26), 5687–5698. 10.1523/JNEUROSCI.2466-20.2021

Ambrus, G. G., Kaiser, D., Cichy, R. M., & Kovács, G. (2019). The Neural Dynamics of Familiar Face Recognition. Cerebral Cortex, 29(11), 4775–4784. 10.1093/cercor/bhz010

Andrews, S., Burton, A. M., Schweinberger, S. R., & Wiese, H. (2017). Event-related potentials reveal the development of stable face representations from natural variability. The Quarterly Journal of Experimental Psychology, 70(8), 1620–1632. 10.1080/17470218.2016.1195851

Bae, G.-Y. (2020). The time course of face representations during perception and working memory maintenance. Cerebral Cortex Communications, 2(1). 10.1093/texcom/tgaa093

Castro-Laguardia, A. M., Ontivero-Ortega, M., Morato, C., Lucas, I., Vila, J., Bobes León, M. A., & Muñoz, P. G. (2024). Familiarity Processing through Faces and Names: Insights from Multivoxel Pattern Analysis. Brain Sciences, 14(1), 39. 10.3390/brainsci14010039

Corriveau, A., Kidder, A., Teichmann, L., Wardle, S. G., & Baker, C. I. (2023). Sustained neural representations of personally familiar people and places during cued recall. Cortex, 158, 71–82. 10.1016/j.cortex.2022.08.014

Dalski, A., Kovács, G., & Ambrus, G. G. (2022). Evidence for a general neural signature of face familiarity. Cerebral Cortex, 32(12), 2590–2601. 10.1093/cercor/bhab366

Dalski, A., Kovács, G., & Ambrus, G. G. (2023). No semantic information is necessary to evoke general neural signatures of face familiarity: Evidence from cross-experiment classification. Brain Structure and Function, 228(2), 449–462. 10.1007/s00429-022-02583-x

Delorme, A. (2023). EEG is better left alone. Scientific Reports, 13(1), 2372. 10.1038/s41598-023-27528-0

Delorme, A., & Makeig, S. (2004). EEGLAB: An open source toolbox for analysis of single-trial EEG dynamics including independent component analysis. Journal of Neuroscience Methods, 134(1), 9–21. 10.1016/j.jneumeth.2003.10.009

Dijkstra, N., Bosch, S. E., & van Gerven, M. A. J. (2019). Shared Neural Mechanisms of Visual Perception and Imagery. Trends in Cognitive Sciences, 23(5), 423–434. 10.1016/j.tics.2019.02.004

Dijkstra, N., Mostert, P., Lange, F. P. de, Bosch, S., & van Gerven, M. A. (2018). Differential temporal dynamics during visual imagery and perception. eLife, 7, e33904. 10.7554/eLife.33904

Dijkstra, N., Zeidman, P., Ondobaka, S., van Gerven, M. a. J., & Friston, K. (2017). Distinct Top-down and Bottom-up Brain Connectivity During Visual Perception and Imagery. Scientific Reports, 7(1), 5677. 10.1038/s41598-017-05888-8

Dobs, K., Isik, L., Pantazis, D., & Kanwisher, N. (2019). How face perception unfolds over time. Nature Communications, 10(1), 1258. 10.1038/s41467-019-09239-1

Dubois, S., Rossion, B., Schiltz, C., Bodart, J. M., Michel, C., Bruyer, R., & Crommelinck, M. (1999). Effect of Familiarity on the Processing of Human Faces. NeuroImage, 9(3), 278–289. 10.1006/nimg.1998.0409

Ely, M. M., Kelsey, C., & Ambrus, G. G. (2026). Shared Neural Codes for Emotion Recognition in Emoji and Human Faces. Psychophysiology, 63(3), e70268. 10.1111/psyp.70268.

Fahrenfort, J. J., van Driel, J., van Gaal, S., & Olivers, C. N. L. (2018). From ERPs to MVPA Using the Amsterdam Decoding and Modeling Toolbox (ADAM). Frontiers in Neuroscience, 12. 10.3389/fnins.2018.00368

Faul, F., Erdfelder, E., Lang, A. G., & Buchner, A. (2007). G*Power 3: A flexible statistical power analysis program for the social, behavioral, and biomedical sciences. Behavior research methods, 39(2), 175–191. 10.3758/BF03193146

Gainotti, G. (2007). Different patterns of famous people recognition disorders in patients with right and left anterior temporal lesions: a systematic review. Neuropsychologia, 45(8), 1591–1607. 10.1016/j.neuropsychologia.2006.12.013

Ganis, G., Thompson, W. L., & Kosslyn, S. M. (2004). Brain areas underlying visual mental imagery and visual perception: An fMRI study. Cognitive Brain Research, 20(2), 226–241. 10.1016/j.cogbrainres.2004.02.012

Gobbini, M. I., & Haxby, J. V. (2006). Neural response to the visual familiarity of faces. Brain Research Bulletin, 71(1), 76–82. 10.1016/j.brainresbull.2006.08.003

Grootswagers, T., Wardle, S. G., & Carlson, T. A. (2017). Decoding Dynamic Brain Patterns from Evoked Responses: A Tutorial on Multivariate Pattern Analysis Applied to Time Series Neuroimaging Data. Journal of Cognitive Neuroscience, 29(4), 677–697. 10.1162/jocn_a_01068

Hayama, H. R., Johnson, J. D., & Rugg, M. D. (2008). The relationship between the right frontal old/new ERP effect and post-retrieval monitoring: Specific or non-specific? Neuropsychologia, 46(5), 1211–1223. 10.1016/j.neuropsychologia.2007.11.021

Hu, Y., & Yu, Q. (2023). Spatiotemporal dynamics of self-generated imagery reveal a reverse cortical hierarchy from cue-induced imagery. bioRxiv. 10.1101/2023.01.25.525474

Ishai, A., Haxby, J. V., & Ungerleider, L. G. (2002). Visual imagery of famous faces: Effects of memory and attention revealed by fMRI. NeuroImage, 17(4), 1729–1741. 10.1006/nimg.2002.1330

Ishai, A., Ungerleider, L. G., & Haxby, J. V. (2000). Distributed Neural Systems for the Generation of Visual Images. Neuron, 28(3), 979–990. 10.1016/S0896-6273(00)00168-9

Kaiser, D., Oosterhof, N. N., & Peelen, M. V. (2016). The Neural Dynamics of Attentional Selection in Natural Scenes. Journal of Neuroscience, 36(41), 10522–10528. 10.1523/JNEUROSCI.1385-16.2016

King, J.-R., & Dehaene, S. (2014). Characterizing the dynamics of mental representations: The temporal generalization method. Trends in Cognitive Sciences, 18(4), 203–210. 10.1016/j.tics.2014.01.002

Klink, H., Kaiser, D., Stecher, R., Ambrus, G. G., & Kovács, G. (2023). Your place or mine? The neural dynamics of personally familiar scene recognition suggests category independent familiarity encoding. Cerebral Cortex, 33(24), 11634–11645. 10.1093/cercor/bhad397

Kovács, G. (2020). Getting to Know Someone: Familiarity, Person Recognition, and Identification in the Human Brain. Journal of Cognitive Neuroscience, 32(12), 2205–2225. 10.1162/jocn_a_01627

Kutsche, J., Howard, C., Drew, W., Michel, M., Cohen, A. L., Fox, M. D., & Kletenik, I. (2025). Visual Mental Imagery and Aphantasia Lesions Map onto a Convergent Brain Network. medRxiv. 10.1101/2025.05.23.25328072

Li, C., Burton, A. M., Ambrus, G. G., & Kovács, G. (2022). A neural measure of the degree of face familiarity. Cortex, 155, 1–12. 10.1016/j.cortex.2022.06.012

Liu, J., Zhan, M., Hajhajate, D., Spagna, A., Dehaene, S., Cohen, L., & Bartolomeo, P. (2025). Visual mental imagery in typical imagers and in aphantasia: A millimeter-scale 7-T fMRI study. Cortex, 185, 113–132. 10.1016/j.cortex.2025.01.013

Maris, E., & Oostenveld, R. (2007). Nonparametric statistical testing of EEG- and MEG-data. Journal of Neuroscience Methods, 164(1), 177–190. 10.1016/j.jneumeth.2007.03.024

O’Craven, K. M., & Kanwisher, N. (2000). Mental imagery of faces and places activates corresponding stimulus-specific brain regions. Journal of Cognitive Neuroscience, 12(6), 1013–1023. 10.1162/08989290051137549

Peirce, J., Gray, J. R., Simpson, S., MacAskill, M., Höchenberger, R., Sogo, H., Kastman, E., & Lindeløv, J. K. (2019). Psychopy2: Experiments in behavior made easy. Behavior Research Methods, 51(1), 195–203. 10.3758/s13428-018-01193-y

Ragni, F., Lingnau, A., & Turella, L. (2021). Decoding category and familiarity information during visual imagery. NeuroImage, 241, 118428. 10.1016/j.neuroimage.2021.118428

Ragni, F., Tucciarelli, R., Andersson, P., & Lingnau, A. (2020). Decoding stimulus identity in occipital, parietal and inferotemporal cortices during visual mental imagery. Cortex, 127, 371–387. 10.1016/j.cortex.2020.02.020

Schweinberger, S. R., Pickering, E. C., Jentzsch, I., Burton, A. M., & Kaufmann, J. M. (2002). Event-related brain potential evidence for a response of inferior temporal cortex to familiar face repetitions. Brain Research. Cognitive Brain Research, 14(3), 398–409. 10.1016/s0926-6410(02)00142-8

Scrivener, C. L., Teed, J. A., & Silson, E. H. (2025). Visual imagery of familiar people and places in category selective cortex. Neuroscience of Consciousness, 2025(1), niaf006. 10.1093/nc/niaf006

Shah, N. J., Marshall, J. C., Zafiris, O., Schwab, A., Zilles, K., Markowitsch, H. J., & Fink, G. R. (2001). The neural correlates of person familiarity. A functional magnetic resonance imaging study with clinical implications. Brain, 124(Pt 4), 804–815. 10.1093/brain/124.4.804

Shatek, S. M., Grootswagers, T., Robinson, A. K., & Carlson, T. A. (2019). Decoding Images in the Mind’s Eye: The Temporal Dynamics of Visual Imagery. Vision, 3(4), 53. 10.3390/vision3040053

Silson, E. H., Steel, A., Kidder, A., Gilmore, A. W., & Baker, C. I. (2019a). Distinct subdivisions of human medial parietal cortex support recollection of people and places. eLife, 8, e47391. 10.7554/eLife.47391

Silson, E. H., Gilmore, A. W., Kalinowski, S. E., Steel, A., Kidder, A., Martin, A., & Baker, C. I. (2019b). A posterior–anterior distinction between scene perception and scene construction in human medial parietal cortex. The Journal of Neuroscience, 39(4), 705–717. 10.1523/JNEUROSCI.1219-18.2018

Spagna, A., Hajhajate, D., Liu, J., & Bartolomeo, P. (2021). Visual mental imagery engages the left fusiform gyrus, but not the early visual cortex: A meta-analysis of neuroimaging evidence. Neuroscience and Biobehavioral Reviews, 122, 201–217. 10.1016/j.neubiorev.2020.12.029

Spagna, A., Heidenry, Z., Miselevich, M., Lambert, C., Eisenstadt, B. E., Tremblay, L., Liu, Z., Liu, J., & Bartolomeo, P. (2024). Visual mental imagery: Evidence for a heterarchical neural architecture. Physics of Life Reviews, 48, 113–131. 10.1016/j.plrev.2023.12.012

Steel, A., Billings, M. M., Silson, E. H., & Robertson, C. E. (2021). A network linking scene perception and spatial memory systems in posterior cerebral cortex. Nature Communications, 12(1), 2632. 10.1038/s41467-021-22848-z

Treder, M. S., Charest, I., Michelmann, S., Martín-Buro, M. C., Roux, F., Carceller-Benito, F., Ugalde-Canitrot, A., Rollings, D. T., Sawlani, V., Chelvarajah, R., Wimber, M., Hanslmayr, S., & Staresina, B. P. (2021). The hippocampus as the switchboard between perception and memory. Proceedings of the National Academy of Sciences, 118(50). 10.1073/pnas.2114171118

Wiese, H., Hobden, G., Siilbek, E., Martignac, V., Flack, T. R., Ritchie, K. L., Young, A. W., & Burton, A. M. (2022). Familiarity is familiarity is familiarity: Event-related brain potentials reveal qualitatively similar representations of personally familiar and famous faces. *Journal of Experimental Psychology: Learning*, Memory, and Cognition, 48(8), 1144– 1164. 10.1037/xlm0001063

Wiese, H., Schweinberger, S. R., & Kovács, G. (2024). The neural dynamics of familiar face recognition. Neuroscience & Biobehavioral Reviews, 167, 105943. 10.1016/j.neubiorev.2024.105943

Wiese, H., Tüttenberg, S. C., Ingram, B. T., Chan, C. Y., Gurbuz, Z., Burton, A. M., & Young, A. W. (2019). A robust neural index of high face familiarity. Psychological Science, 30(2), 261–272. 10.1177/0956797618813572

Wilding, E. L., & Rugg, M. D. (1996). An event-related potential study of recognition memory with and without retrieval of source. Brain, 119 ( *Pt 3*), 889–905. 10.1093/brain/119.3.889

Xie, S., Kaiser, D., & Cichy, R. M. (2020). Visual Imagery and Perception Share Neural Representations in the Alpha Frequency Band. Current Biology, 30(13), 2621–2627.e5. 10.1016/j.cub.2020.04.074

Zhang, G., & Luck, S. J. (2025). Assessing the impact of artifact correction and artifact rejection on the performance of SVM-and LDA-based decoding of EEG signals. NeuroImage, 316, 121304. 10.1016/j.neuroimage.2025.121304

