## Supplementary Materials for "The neural dynamics of familiarity representation during cued recall"

**Supplementary Information 1. Pilot study to validate familiarity of symbols and unfamiliar faces**

A pilot study was conducted to validate the familiarity categorization of the selected symbols, used in the main experiment. 5 exemplars of familiar (Pirates of the Caribbean symbol corresponding to Johnny Depp, Iron Man symbol corresponding to Robert Downey Jr., Harry Potter symbol corresponding to Emma Watson, United States of America flag/symbols corresponding to Donald Trump, CDU symbol corresponding to Angela Merkel) and unfamiliar (Infogames, Macmillan, Trabzonspor, Mahindra, Meralco) symbols were presented and participants were asked to imagine a face they associated with the symbol shown. The name of the person they imagined was asked, along with how vividly they imagined the face (Supplementary Fig. 1; Likert ratings with 1 corresponding to not vivid at all and 4 corresponding to very vivid). We also asked if the participants have ever encountered the symbol in the end (in order to make sure the unfamiliar symbols were indeed unfamiliar even if the participants could not name them). We calculated the recognition accuracy (Supplementary Fig. 2) as the proportion of correct answers (whether the imagined face corresponds to someone actually related to the symbol; if multiple participants are related to a symbol, we accepted all possible answers as correct) across all trials for a given symbol, per each subject. Then we aggregated the accuracy scores across subjects. The results validated our stimulus selection in the sense that none of the subjects were able to recognize the unfamiliar symbols, and all the familiar symbols had a recognition accuracy above 50%.  Although we initially planned to use Emma Watson as the familiar identity associated with the Harry Potter symbol set, most participants responded with “Harry Potter”. Based on this outcome and to align the pilot with the final experiment, we ultimately used Daniel Radcliffe's face (actor playing the character Harry Potter) as the familiar identity in the main study. To keep the identity set consistent, we also removed Angela Merkel so that all identities were male, and we removed Meralco to ensure a matched number of familiar and unfamiliar identities in the final stimulus set.

To validate that the identities we would use for the experiment were indeed unfamiliar to a German audience, participants were also shown face images of 5 celebrities that were famous outside Germany (Leandra Leal, Goran Bogdan, Jorge Drexler, Alban Skënderaj, Uğurkan Erez). None of the participants could recognize any of the unfamiliar identities we presented. To match the number of familiar identities and to keep only males, Leandra Leal was removed from unfamiliar identities in the main experiment.

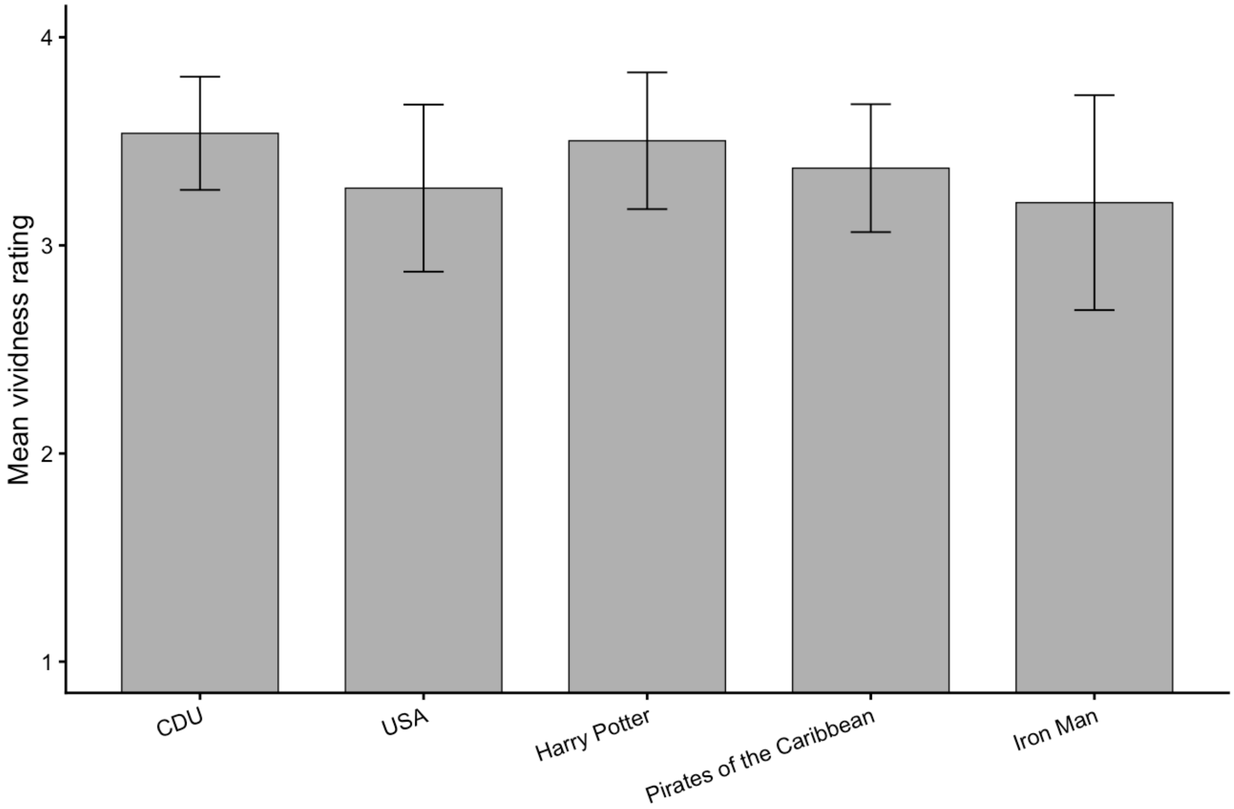

**Supplementary Figure 1**. *Pilot vividness ratings for correct identifications. Mean vividness ratings (1-4) for each symbol, computed using correct trials only (Score = 1). Bars show subject-level mean vividness per symbol averaged across participants; error bars indicate 95% confidence intervals across subjects.*

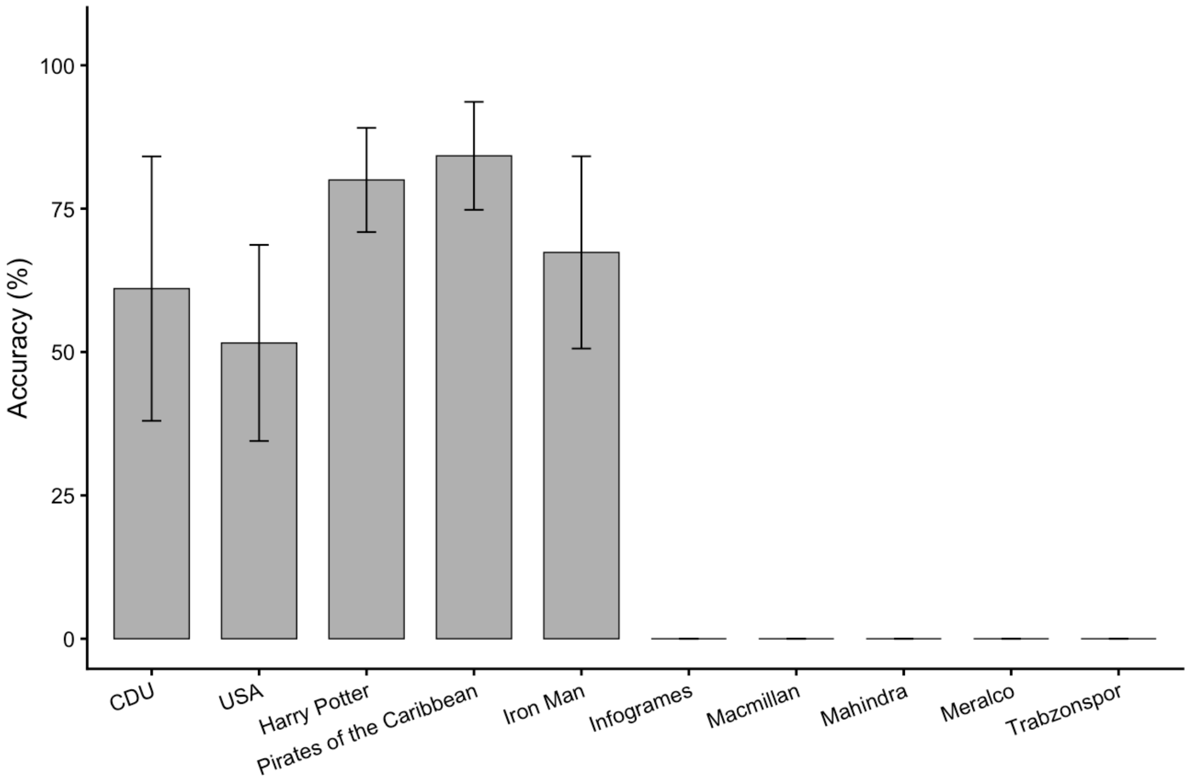


**Supplementary Figure 2**. *Pilot validation of symbol familiarity. Recognition accuracy (% correct) for each symbol. Bars show subject-level mean accuracy (each participant's proportion correct for a given symbol, averaged across subjects); error bars indicate 95% confidence intervals.*

**Supplementary Information 2. Post-cued recall behavioral evaluation**

After the cued recall phase, participants were given a handout and were asked to write the names of the people they imagined for each of the symbols shown. We took the answers that were the same with the faces shown on the localizer phase as correct (Supplementary Fig. 3).

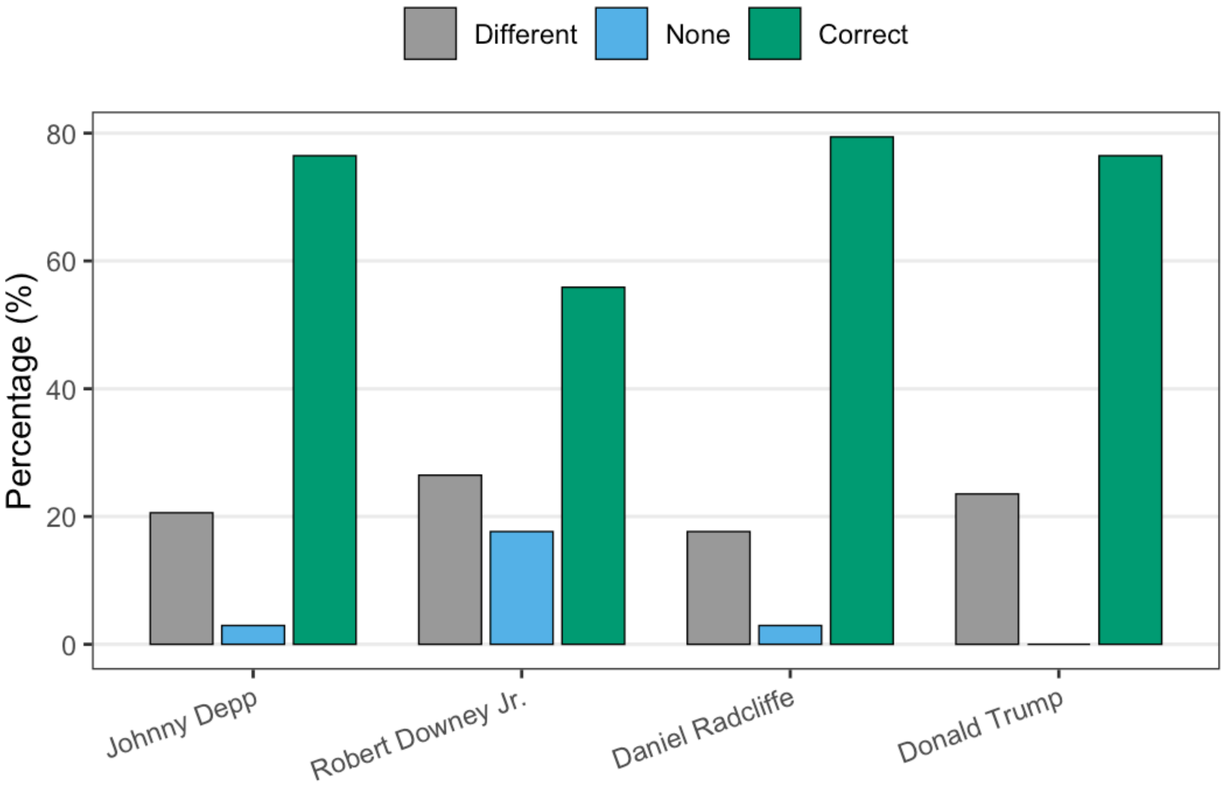


**Supplementary Figure 3**. *Percentage of responses in each accuracy category (Different, None, Correct) for the symbols associated with four familiar identities (Johnny Depp, Robert Downey Jr., Daniel Radcliffe, Donald Trump) in the post-cued recall behavioral evaluation. Bars show the proportion of post-task responses assigned to each category for each identity, displayed as side-by-side bars. Colors indicate response category (grey = Different, blue = None, green = Correct).*

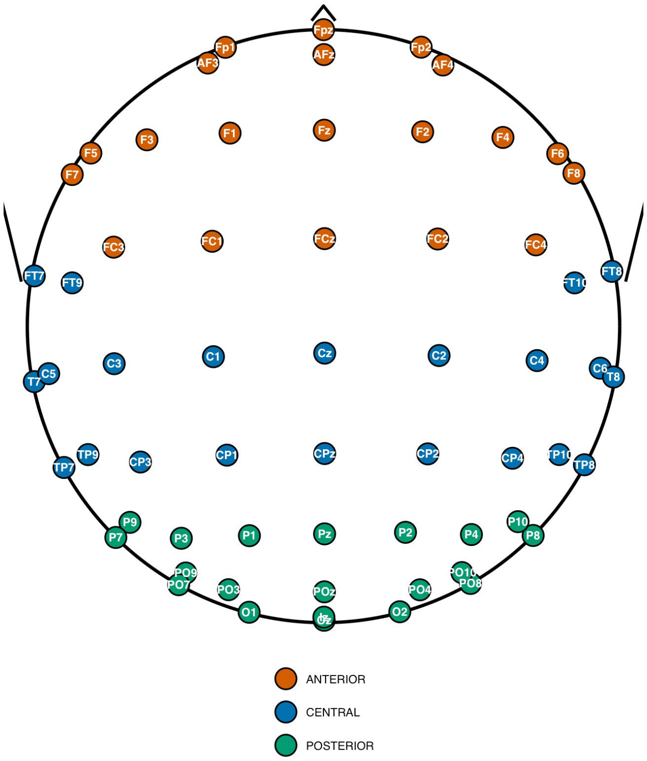


**Supplementary Figure 4.** *Electrode groups used for the ROI-based decoding analyses. Scalp schematic showing the electrodes included in the left and right anterior, central, and posterior ROIs. Midline electrodes were included in both the corresponding left and right ROI groups.*

**Supplementary Information 3. Behavioral validation of the cued recall task**

Supplementary Fig. 5 depicts the behavioral validation analyses reported in the main manuscript, including the difference in vividness ratings between familiar and unfamiliar cued-recall trials and the relationship between vividness and congruent/intended person recall.

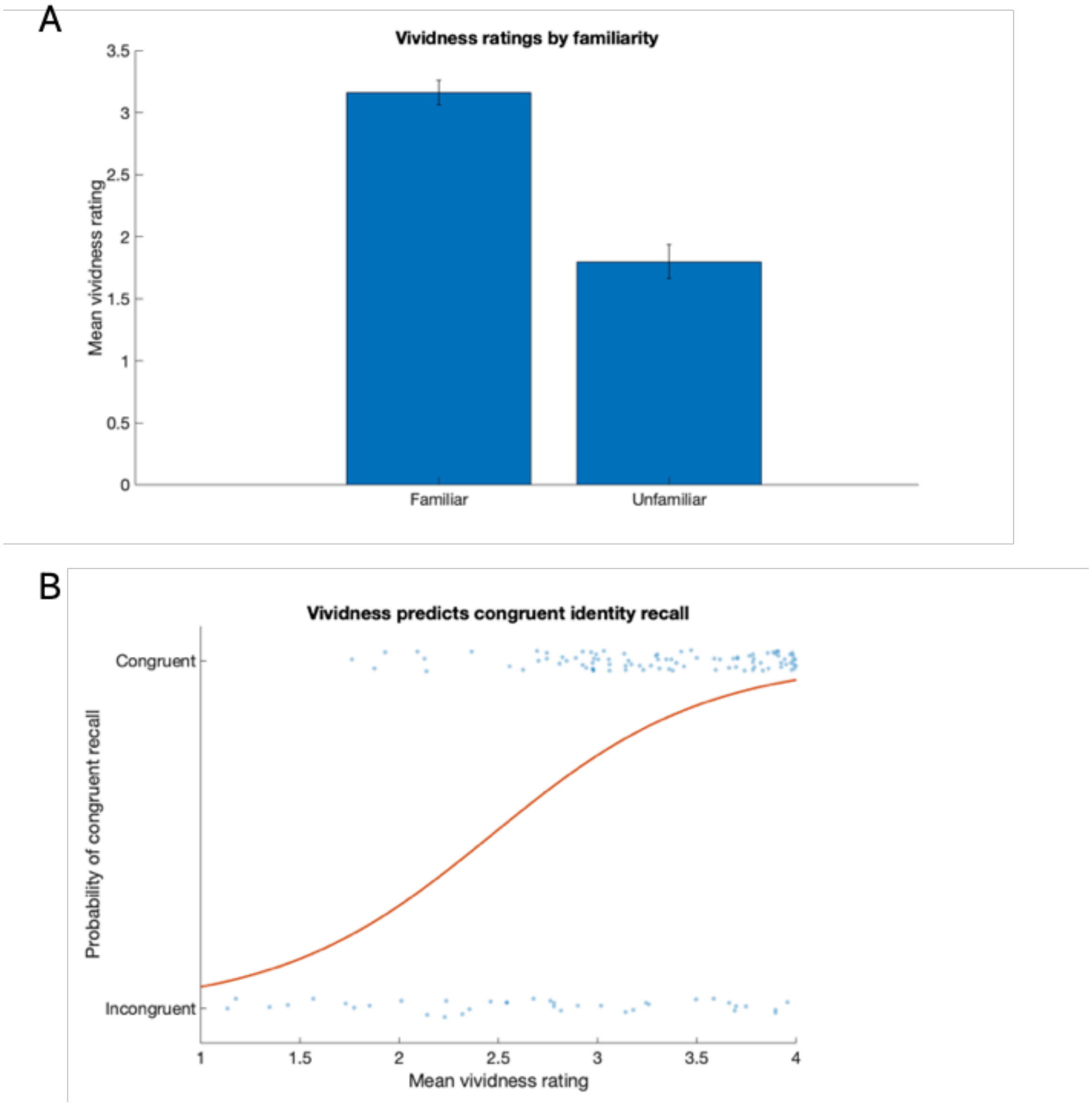


**Supplementary Figure 5.** *Behavioral validation of the cued recall task.* ***(A)*** *Mean vividness ratings for familiar and unfamiliar symbol trials. Error bars indicate the standard error of the mean.* ***(B)*** *Predicted probability (red line) of congruent/intended person recall as a function of mean vividness rating based on the mixed-effects logistic regression. Points represent participant-by-identity observations.*

**Supplementary Information 4. Robustness analysis excluding participants with frequent reported person imagery for unfamiliar symbols**

To examine whether cross-domain generalization was influenced by participants who frequently imagined people in response to unfamiliar symbols, we repeated the cross-domain temporal generalization analysis after excluding participants who reported imagining a person for more than two unfamiliar symbols. Seven participants met this criterion, resulting in a reduced sample of 27 participants.

Cross-domain temporal generalization analysis with the reduced sample has shown that the whole-brain significant cluster was preserved (train: 60-1190 ms; test: 190-4590 ms) along with the other significant bilateral early posterior clusters (left: train 190-1190 ms, test 100-1150 ms; right: train 250-940 ms, test 240-1200 ms), a later cluster in left posterior (train: 420-1190 ms; test: 1190-2760 ms) and a prolonged late cluster in right anterior (train: 590-1190 ms; test: 1890-4260 ms). In the end, the general pattern of cross-domain generalization remained comparable to that observed in our full sample (Supplementary Fig. 6).

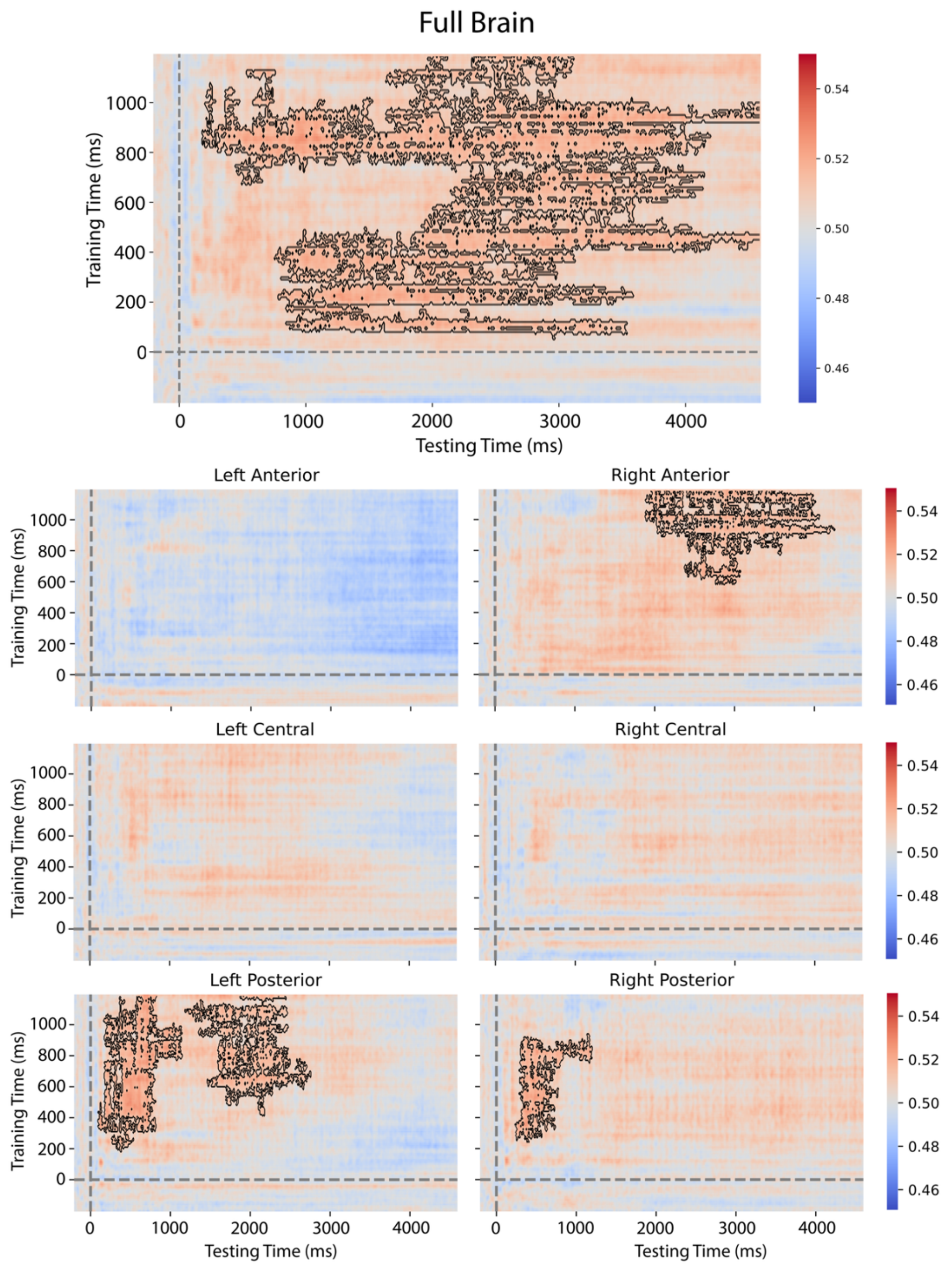


**Supplementary Figure 6.** *Cross-domain temporal generalization of familiarity in the reduced sample (N = 27). Horizontal rows indicate timepoints that the classifiers were trained on (perceptual data of the localizer trials) and vertical columns indicate timepoints that the classifier was tested (data from the cued recall trials) on. Seven participants, who reported imagining a person for more than two unfamiliar symbols were excluded from this analysis. Decoding accuracy*

*is plotted as a proportion; 0.50 corresponds to 50% accuracy and to chance level performance. Significant temporal clusters are surrounded with black lines. Significance was tested by 10000 one-tailed cluster-based permutations, p< 0.05.*

**Supplementary Information 5. Reverse train-test direction cross-domain temporal generalization analysis**

The complementary reverse direction cross-domain analysis, in which classifiers were trained on the cued-recall data and tested on the localizer (face perception) data, revealed significant temporal generalization in the bilateral posterior and right anterior ROIs (Supplementary Fig. 7), no significant whole-brain cluster was observed in the reverse direction. The posterior ROIs showed generalization from relatively early cued recall training periods to perceptual familiarity processing (left: train 320-1170 ms, test 180-1190 ms; right: train 250-870 ms, test 220-1100 ms), whereas the right anterior ROI showed significant generalization from later cued recall training periods (train: 1460-2340 ms; test: 360-1190 ms). Overall, the reverse analysis therefore showed a broadly convergent, although not identical, ROI level pattern to the primary perception-to-recall direction analysis.

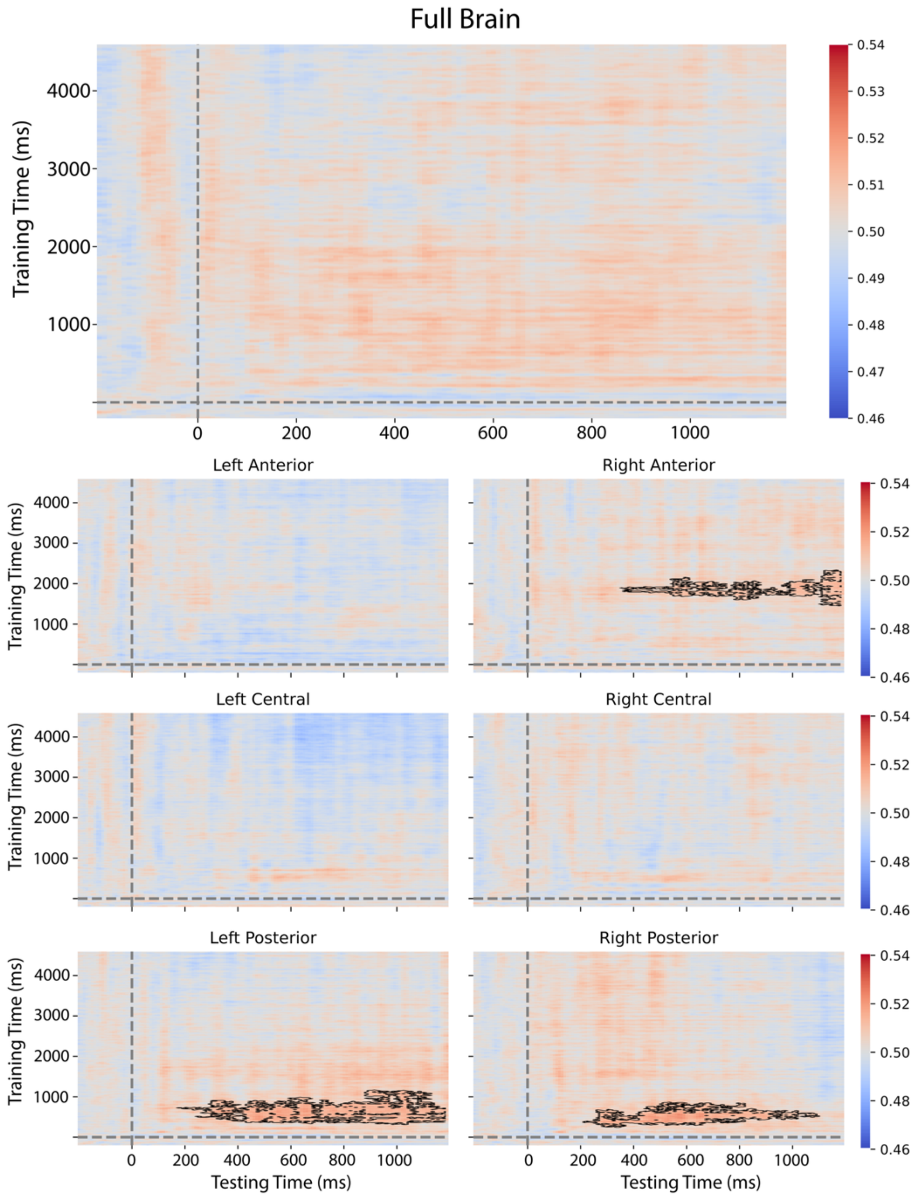


**Supplementary Figure 7.** *Reverse direction cross-domain temporal generalization of familiarity. Horizontal rows indicate timepoints that the classifiers were trained on (data from cued recall trials) and vertical columns indicate timepoints that the classifier was tested (perceptual face data from localizer trials) on. This analysis represents the complementary reverse direction of the primary perception-to-recall cross-domain analysis. Decoding accuracy is plotted as a proportion; 0.50 corresponds to 50% accuracy and to chance level performance. Significant*

*temporal clusters are surrounded with black lines. Significance was tested by 10000 one-tailed cluster-based permutations, p< 0.05.*

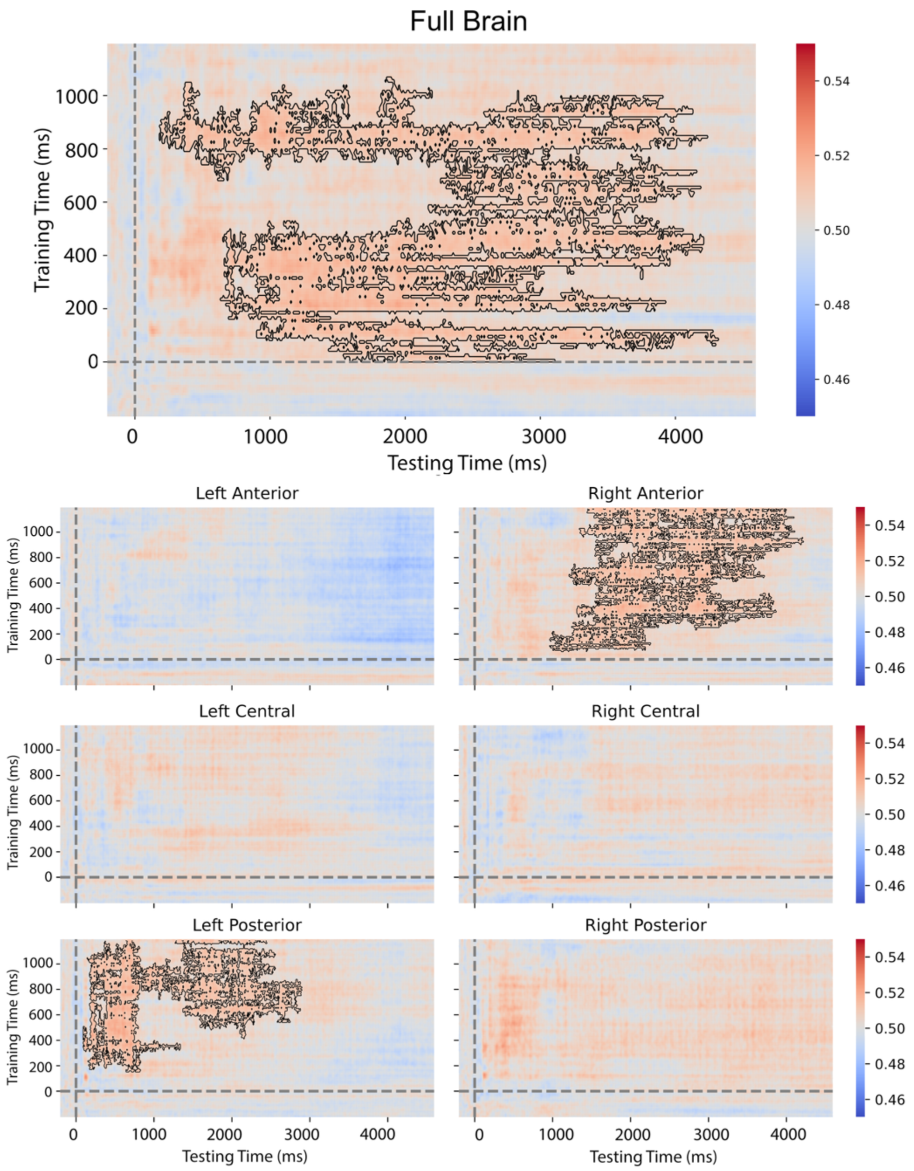


**Supplementary Figure 8.** *Cross-domain decoding analysis with the alternative pointwise threshold of p<0.1. Decoding accuracy is plotted as a proportion; 0.50 corresponds to 50% accuracy and to chance level performance. Significant temporal clusters are surrounded with black lines. Significance was tested by 10000 one-tailed cluster-based permutations, p< 0.05.*
